# Integrative Genome Modeling Platform reveals essentiality of rare contact events in 3D genome organizations

**DOI:** 10.1101/2021.08.22.457288

**Authors:** Lorenzo Boninsegna, Asli Yildirim, Guido Polles, Sofia A. Quinodoz, Elizabeth Finn, Mitchell Guttman, Xianghong Jasmine Zhou, Frank Alber

**Affiliations:** Institute of Quantitative and Computational Biosciences (QCBio), University of California Los Angeles, Los Angeles, CA 90095, USA; Department of Microbiology, Immunology, and Molecular Genetics, University of California Los Angeles, 520 Boyer Hall, Los Angeles, CA 90095, USA; Division of Biology and Biological Engineering, California Institute of Technology, Pasadena, CA 91125, USA; National Cancer Institute, NIH, Bethesda, MD 20892, USA; Department of Pathology, David Geffen School of Medicine, University of California Los Angeles, 10833 Le Conte Ave, Los Angeles, CA 90095, USA

## Abstract

A multitude of sequencing-based and microscopy technologies provide the means to unravel the relationship between the three-dimensional (3D) organization of genomes and key regulatory processes of genome function. However, it remains a major challenge to systematically integrate all available data sources to characterize the nuclear organization of genomes across different spatial scales. Here, we develop a multi-modal data integration approach to produce genome structures that are highly predictive for nuclear locations of genes and nuclear bodies, local chromatin compaction, and spatial segregation of functionally related chromatin.

By performing a quantitative assessment of the predictive power of genome structures generated from different data combinations, we demonstrate that multimodal data integration can compensate for systematic errors and missing values in some of the data and thus, greatly increases accuracy and coverage of genome structure models. We also show that alternative combinations of different orthogonal data sources can converge to models with similar predictive power. Moreover, our study reveals the key contributions of low-frequency inter-chromosomal contacts (e.g., “rare” contact events) to accurately predicting the global nuclear architecture, including the positioning of genes and chromosomes. Overall, our results highlight the benefits of multi-modal data integration for genome structure analysis, available through the Integrative Genome structure Modeling (IGM) software package that we introduce here.

## Introduction

The spatial organization of eukaryotic genomes plays crucial roles in regulation of transcription, replication and cell differentiation, while malfunctions in chromatin structure is linked to disease, including cancer and premature aging disorders^1,2^. Advances in chromosome conformation capture (3C)-based^3–10^ and ligation-free methods^11–13^ and, most recently, live-cell and superresolution microscopy^14–18^, have shed light into key elements of genome structure organization, including the genome-wide detection of chromatin loops^19,20^, topological associating domains (TADs)^21^ that modulate long-range promoter-enhancer interactions^12,22^ as well as the segregation of chromatin into nuclear compartments^8,10,23–26^. Each technology probes different aspects of genome architecture at different resolutions^1,27–29^.

These complementary methods provide a renewed opportunity to generate quantitative, highly predictive structural models of the entire nuclear organization^30^. Embedding data into three dimensional structures is beneficial for a variety of reasons. First, all data itself originates from (often a large population of) 3D structures; so, reverse engineering that data and relating it back to an ensemble of representative 3D structures must be possible and appears to be the natural way for integrating data from complementary methods via an appropriate representation of experimental errors and uncertainties. Second, generating structures consistent with multi-modal data from heterogeneous and independent sources allows cross-validation of orthogonal data itself. Finally, 3D structures give access to features that are not immediately visible in the original input data set, which can be compared with experimental data tailored to assess model predictivity. Yet, embedding data into 3D structures is a challenging task: not only is there no established protocol for data interpretation and modeling, but genome structures are dynamic in nature and can substantially vary between individual cells. A probabilistic description is thus needed surpassing traditional structural modeling that limits to a single equilibrium structure, or a small number of metastable structures.

There are several data-driven and mechanistic modeling strategies, which differ in the functional interpretation of data and sampling strategies, for generating an ensemble of 3D genome structures statistically consistent with it^23,25,26,31–51^. These 3D structures are then examined to derive structure-function correlations and make quantitative predictions about structural features of genomic regions, study their cell-to-cell variabilities and link these to functional observations. Most strategies have relied primarily on Hi-C data, which is abundant and straightforward to interpret in terms of chromatin contacts. However, data from a single experimental method cannot possibly capture all aspects of the spatial genome organization. Integrating data from a wide range of technologies, each with complementary strengths and limitations, will likely increase accuracy and coverage of genome structure models. Several methods were adapted to combine Hi-C with one other data source, such as lamin ChIP-seq data (Chrom3D)^52^, lamina DamID^38^, 3D FISH distances^51,53^, super-resolution OligoSTORM/Oligo-DNA-PAINT images^14^ and GPSeq data^54^. Nevertheless, developing hybrid methods that can systematically integrate data from many different technologies to generate structural maps of entire diploid genomes remains a major challenge.

Here we present a population-based deconvolution method that provides an ideal probabilistic framework for comprehensive and multi-modal data integration. Our population-based approach^30,37,45^ de-multiplexes ensemble data into a population of 3D structures, each governed by a unique pseudo-energy function, representing a subset of the data. Therefore, our approach explicitly factors in the heterogeneity of structural features across different cells. The method produces highly predictive models of the folded states of complete diploid genomes, which are statistically consistent with all input data^30^. Thus, our method is distinct from resampling methods^32,42,46,47,55,56^, which generate genome models by multiple sampling of the same scoring function, derived from ensemble data.

Our generalized framework generates fully diploid genome models from integration of four orthogonal data types, ensemble Hi-C^10^, lamina-B1 DamID^24,57,58^, large-scale HIPMap 3D FISH imaging^59,60^ and data from Split-Pool Recognition of Interactions by Tag Extension (SPRITE) experiments^11^. Such models have highly predictive values, and are capable to successfully predict with good accuracy orthogonal experimental data from a variety of other genomics-based and super resolution imaging experiments, such as data from SON-TSA-seq experiments^61^ and DNA-MERFISH imaging^17^. Specifically, our structures predict with good accuracy gene distances to nuclear speckles, gene distances to the nuclear lamina and therefore allow an in-depth analysis of the nuclear microenvironment of genes at a genome-wide scale.

We further demonstrate that integration of multiple data modalities maximizes prediction accuracy and we propose the best combinations of data to maximize genome structure accuracy. For instance, our results highlight that relatively low frequent inter-chromosomal contacts are essential to correctly predict whole genome structure organizations: indeed, a modified Hi-C data set with artificially underrepresented inter-chromosomal contacts severely fails at reproducing the correct global genome architecture. However, integrating additional data sources from other experiments can overcome and compensate for these biases and generates structure populations with still high predictivity accuracy. Our method is generally applicable to any cell type and organisms, with any combination of data available, discussed here.

To the best of our knowledge, our work represents the first effort at integrating orthogonal data types from Hi-C, lamina DamID, 3D HIPMap FISH and DNA SPRITE experiments to produce highly predictive genome structure populations, which ultimately showcases the benefits of multimodal data integration in the context of whole genome modeling. Due to its modular architecture, the method we propose can be easily adapted to incorporate other data types in the modeling pipeline, as we strive for even more realistic and predictive structures to dissect the genome structure-function relationship.

## Results

### Multi-modal data-driven population modeling as an optimization problem

We expand our previous genome modeling framework^37,38,45^ and introduce a generalized formulation for the integration of a variety of orthogonal data to generate a population of full genome structures that simultaneously recapitulate all the data. Our method incorporates univariate, bivariate *and* multivariate data types (**Fig. 1**). Univariate data describes probabilities of individual genomic loci: for example, lamina-B1 DamID data provides univariate information about the contact frequencies of individual loci to the nuclear envelope, while HIPMap 3D FISH provides distributions of radial positions for individual loci. Bivariate data expresses probabilities of pairs of loci, such as contact frequencies between two chromatin regions from Hi-C data or distance distributions between pairs of loci from HIPMap 3D FISH experiments. Multivariate data represent information tied to more than two loci, for instance the spatial co-location of multiple chromatin regions in single cells as detected in SPRITE experiments. Our method incorporates both ensemble and single cell data by deconvoluting ensemble data into a population of distinct single-cell genome structures, which cumulatively recapitulate all input information. Our model is defined as a population of *S* diploid genome structures ***X*** = {***X***_1_, ***X***_2_,…***X***_*S*_}, where each structure ***X***_*s*_ is represented by a set of 3D vectors representing the coordinates of all diploid chromatin regions. Given univariate (*U*), bivariate (***M***) and multivariate (***T***) input data, we aim to estimate the structure population 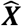 such that the likelihood *P*(*U, **M, T|X***) is maximized, i.e.

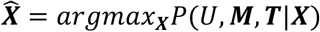

**Figure 1.**
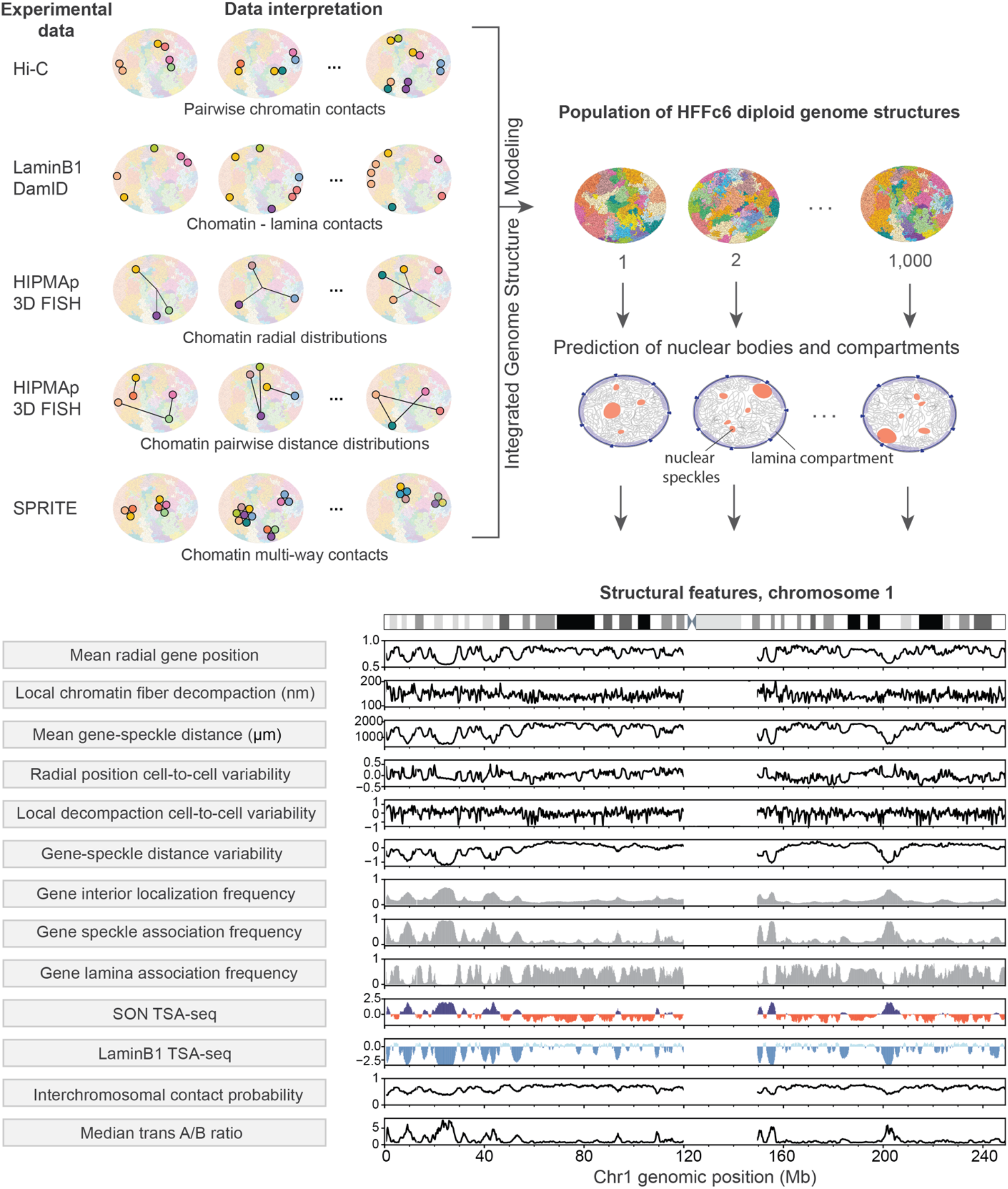
Prediction of the nuclear microenvironments of genes from genome structures. (Top) Schematic view of the data-driven modeling approach. Information provided by orthogonal data modalities (Hi-C, lamina DamID, radial and pairwise HIPMap 3D FISH and DNA SPRITE) are used as input to the Integrative Genome Structure Modeling (IGM) platform to generate a population of *S* = 1000 diploid genome structures. Structures can be used to predict locations of nuclear bodies and compartments (nuclear speckles and lamina compartment), which can serve as reference points to describe locations of genes and the genome architecture. (Bottom) The predicted genome structure population gives access to a large number of structural features (listed on the left), which collectively describe the nuclear microenvironment of genes on a genome-wide scale.

Because most experiments, such as Hi-C and Lamina DamID, provide data that are averaged over a large population of cells, and often produce unphased data, they do not reveal which contacts co-exist in which structure or between which homologue chromosome copies. To represent this missing information at single cell and diploid level, we introduce indicator tensors ***V, W, R*** as latent variables that augment all missing information in ***U, M, T***, respectively. We therefore jointly optimize all latent variables and genome structures using a variant of the Expectation-maximization (EM) method that iteratively optimizes local approximations of the log likelihood function.

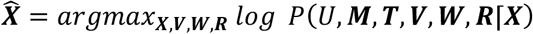

The solution of the high-dimensional maximum likelihood problem requires extensive exploration of the space of all genome structure populations. This problem is solved by an iterative optimization procedure using a series of optimization strategies for efficient and scalable model estimation (*Methods*, **Extended Data Figure 1**)^37,38,45,62^. Convergence to an optimal population 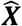 is reached when the models statistically reproduce all the input data. Details on the implementation, optimization parameters and preprocessing of each experimental data source are provided in the *Methods* and *Supplementary Information*. The optimized structure population 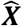 is then used to determine locations of nuclear bodies in each cell model, which in turn serve as reference points to calculate a host of structural features. These features allow a thorough characterization of the nuclear microenvironment of each gene^30^ (**Fig. 1**).

### Comprehensive data-driven genome population structures of HFFc6 cell line

To showcase our data-integration platform we generate a population of 1000 3D diploid genome structures of prolate ellipsoidal HFFc6 fibroblast cell nuclei at 200k base-pair resolution by integrating data from *in situ* Hi-C^63^, lamina-B1 DamID^64^, HIPMap large-scale 3D FISH imaging^59^ and DNA SPRITE experiments^11^. These structures are statistically consistent with all input data: (i) genome-wide Hi-C contact probabilities (Pearson’s correlation of 0.98, **Fig. 2a,b**), (ii) chromatin contact probabilities to the nuclear envelope from lamina B1 DamID experiments (Pearson’s correlation of 0.93, **Fig. 2c,d**), (iii) pairwise distance distributions for 51 pairs of loci from 3D HIPMap experiments (Pearson’s correlation of 1.0 of cross Wasserstein Distances **Fig. 2e,f**) and (iv) chromatin colocalizations for more than 6600 chromatin clusters from SPRITE experiments (**Fig. 2g** and **Extended Data Figure 2d**). Moreover, all our results are highly reproducible in independent replicate optimizations. The full optimization statistics are detailed in **Extended Data Fig. 2b,c,d**.

**Figure 2.**
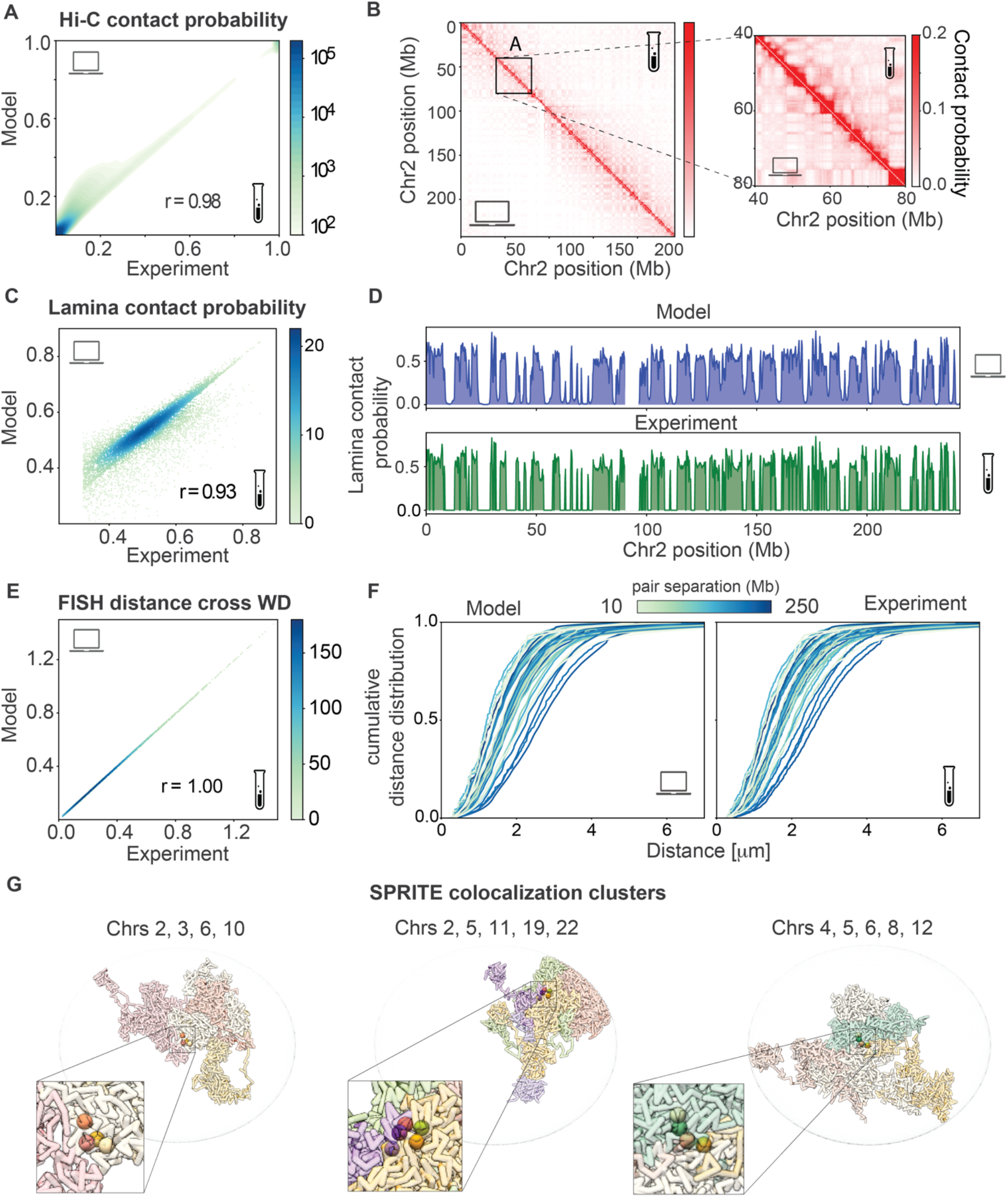
Input data are recapitulated in the genome structure population. (a) Genomewide correlation of Hi-C contact frequencies (inter and intra-chromosomal) between experiment^63^ and simulation (Pearson 0.98). (b) Comparison between experimental (upper diagonal) and simulated (lower diagonal) contact frequency maps for chromosome 2 (left) and zoomed in region (right). (c and d) Correlation of lamin-B1 DamID derived contact probabilities between experiment^64^ and models (c) of genome-wide (Pearson 0.93) and (d) for chromosome 2. (e) Correlation of cross Wasserstein Distances (WD) between experimental FISH data and predictions (Pearson = 1.00) (*Methods*). (f) Cumulative distributions of pairwise FISH distances for the set of 51 pairs of loci measured in FISH experiments^59^, plotted for both models (left) and experiment (right): Colors indicate the sequence separation in the chromosome between imaged loci pairs, with darker hues for larger sequence separations. (g) Typical examples of SPRITE clusters, showing colocalization of loci in a single cell structure: colors distinguish chromosomes, homologues are shown in the same color. Loci in the same SPRITE cluster are also shown enlarged; left cluster: chr2:150,927,500, chr3:6,265,500, chr6:93,928,500, chr10:11,602,500, center cluster: chr2:4,872,500, chr5:23,208,500, chr11:57,966,500, chr19:51,314,500, chr20:42,294,500, right cluster: chr4:42,821,500, chr5:68,438,500, chr6:106,123,500, chr8:85,891,500, chr12:99,185,500. Clusters assayed experimentally^11^ are reproduced in our structures.

We first analyze the predictive value of our models, by simulating orthogonal data from the structures. Specifically, we predict the locations of nuclear speckles in our models, following a previously described procedure^30^. We noticed that chromatin with the 5% lowest average radial positions have generally a high propensity to be associated with nuclear speckles in most cells. Therefore, we identify in each cell model, spatial partitions formed by these chromatin regions. These partitions are detected as highly connected subgraphs in the corresponding chromatin interaction network of each model, and define volume areas, in which these chromatin regions congregate in 3D space (*Methods*). On average the targeted chromatin is fragmented into about 32 spatial partitions per model. We previously discovered that the geometric centers of these spatial partitions serve as excellent approximations of nuclear speckle locations^30^.

SON TSA-seq is an experimental mapping method that determines, on a genome-wide scale, the median distances between any chromatin region and nuclear speckles^61^. To assess the quality of our predicted speckle locations, we simulate the experimental SON TSA-seq process from the folded genome models and predicted speckle locations (*Methods* and **Fig. 1**). The SON-TSA-seq data predicted form our models agrees remarkably well with experiment^65^ (Pearson 0.83, **Fig. 3a**). Moreover, our models confirm the previously described relationship between a chromatin region’s experimental SON TSA-seq value and its mean distance to the nearest speckle^61^.

**Figure 3.**
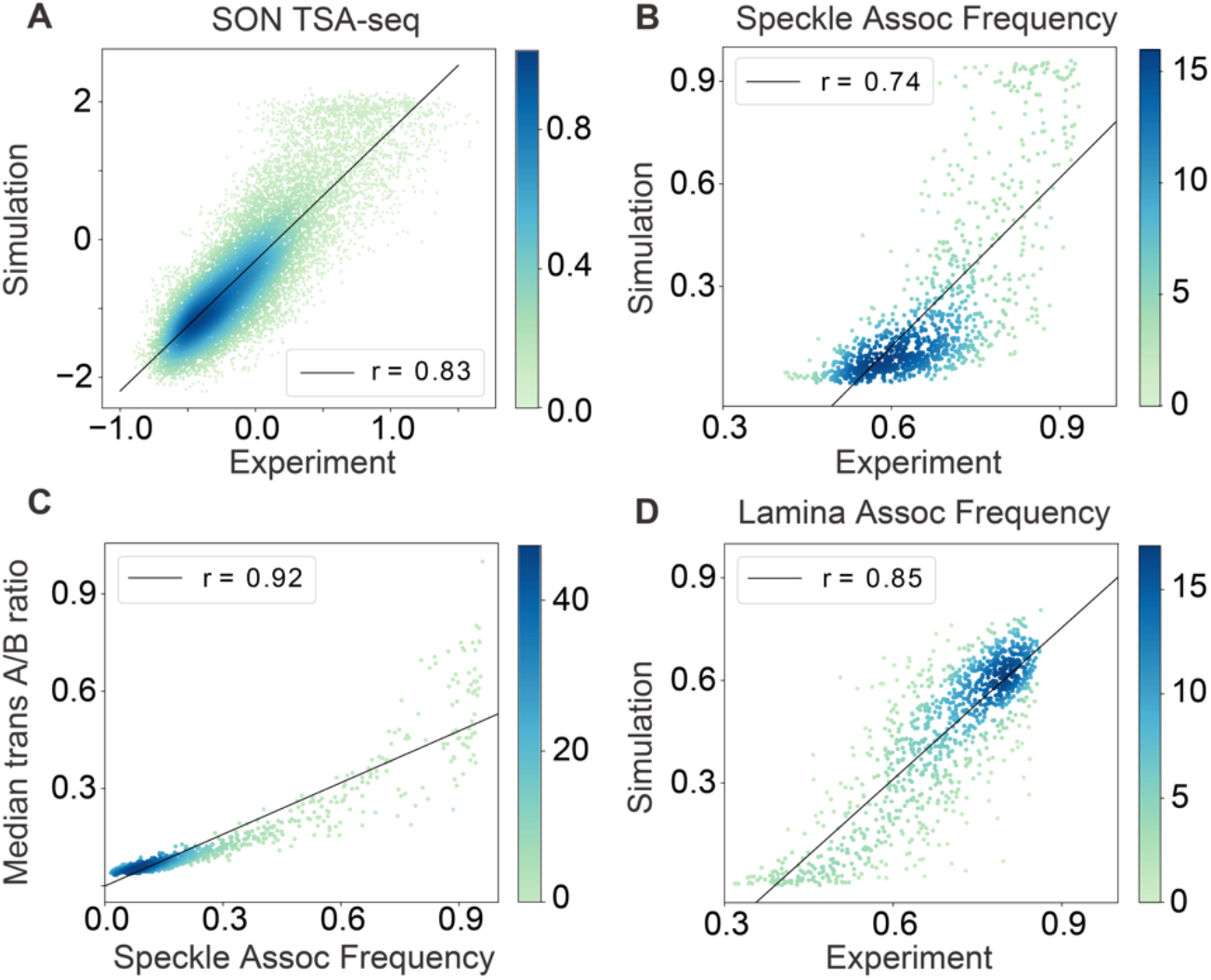
Genome structure population (from HDSF setup) correctly predicts a number of orthogonal experimental observables. (a) Correlation between experimental^65^ and predicted SON TSA-seq data (b) Correlation between predicted and experimental Speckle Associated Frequency (SAF) from DNA-MERFISH imaging; (c) experimentally observed correlation between SAF and transA/B ratio from DNA-MERFISH imaging is nicely reproduced in our genome structures with high correlation. (d) Correlation between experimental Lamina Associated Frequency (LAF) from DNA-MERFISH imaging with predictions from our genome structure population. All scatter plots are colored according to the local density of points, and the Pearson correlation scores are annotated. TSA-seq correlations are genome wide, DNA-MERFISH data correlations involve the 1041 loci studied in the experiment by Su et al^17^.

We then use the predicted speckle locations to determine a gene’s speckle association frequency (SAF), which is defined as the fraction of models, in which a chromatin region is in spatial association to a speckle (*Methods*, **Fig. 1**). A recent super-resolution microscopy study detected the same quantity for approximately 1000 loci by DNA-MERFISH imaging in a related IMR90 cell type^17^. The SAF prediction for these loci from our models shows excellent agreement with the experiments (Pearson correlation 0.71, **Fig. 3b**). Also, Su *et al*. detected the transcript frequency of each imaged loci: interestingly, their transcript frequency shows significant correlation with our simulated SAF (Pearson correlation 0.46), confirming a previous study^30^.

Moreover, we predict for each chromatin region the median trans A/B ratio (*Methods*), defined as the ratio of A and B compartment chromatin forming inter-chromosomal interactions with the target loci. Predicted trans A/B ratios show good agreement with those determined by DNA-MERFISH experiments (Pearson correlation 0.66) and a strong correlation with the SAF (Pearson correlation 0.92, **Fig. 3c**), again confirming previous findings^17,30^.

The lamina associated repressive chromatin compartment is usually located at the nuclear envelope (NE). Thus, we use the location of the NE as a reference point to simulate lamina-B1 TSA-seq data (*Methods*), which measures the mean distances of genomic regions to the nuclear lamina^61^. Moreover, we also calculate the lamina association frequency (LAF) for each genomic region (**Fig. 1**), which also shows excellent agreement with the LAF determined by super resolution DNA-MERFISH imaging^17^ (Pearson 0.84 for LAF, **Fig. 3d**). We also observe an inverse correlation between LAF and SAF (Pearson −0.77), confirming experimental observations.

Overall, the accurate prediction of orthogonal observables assayed in independent experiments highlights the predictive power of our genome structures.

In addition, we also calculate for each chromatin region several other structural features (**Fig. 1** and *Methods*), namely: (i) a chromatin region’s average radial position in the nucleus, (ii) the variability of its radial positions between cells, (iii) the interior localization probability (ILF), defined as the fraction of cells in which a gene is located in the interior region of the nucleus, (iv) the inter chromosomal contact probability, defined as the fraction of interactions formed to other chromosomes as well as (v) the average local chromatin decompaction of the chromatin fiber and its (vi) variability across the population of models. Each chromatin region is therefore defined by a total of 13 structural features, which as a whole define the structural microenvironment of each gene. We have previously^30^ shown that a gene’s microenvironment provides information about its functional potential, in terms of transcription, replication and nuclear compartmentalization.

### Multi-modal data integration improves predictive power

We now investigate how different combinations of data influence model accuracy. We generate four genome populations, each with different combinations of experimental data, and assess their accuracy by comparing predicted SON-TSAseq data, lamina DamID data, SAF, LAF and median trans A/B ratios with those available from experiments (see *Methods*) (**Fig. 4**). For reference, we also assess a population of random chromosome territories constrained within the nuclear volume (*rand*).

**Figure 4.**
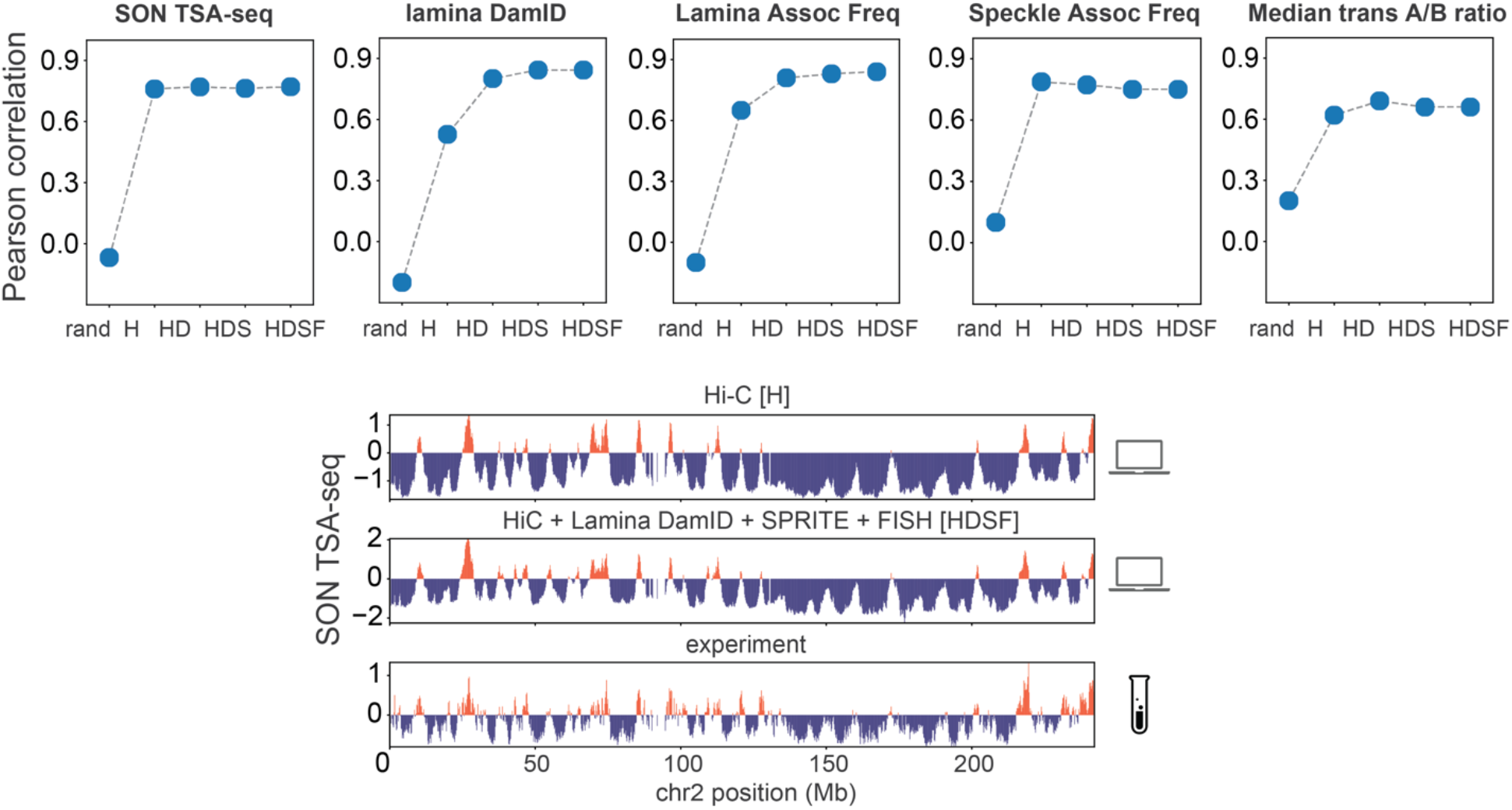
Predictive power and assessment of genome structures increases with integration of more data modalities. (Top) Model accuracy for five different genome structure populations generated from different combinations of experimental input data sets: random chromosome territory (rand), (H) Hi-C only, (HD) Hi-C + lamina DamID data, (HDS) Hi-C + lamina DamID + SPRITE data and (HDSF) Hi-C + lamina DamID + SPRITE + FISH data. (Top, left two panels) Genome-wide Pearson’s correlation coefficients between model predictions and experimental data for experimental SON TSA-seq data, laminB1 DamID. (Top, right three panels) Pearson correlation between experimental and predicted data for Lamina association frequency (LAF), Speckle association frequency (SAF) and trans A/B ratio for 1041 imaged loci from DNA-MERFISH imaging experiments^17^. (Bottom) Comparison between experimental^65^ and predicted SON TSA-seq profile of chromosomes 2, top and bottom respectively. Predicted profiles are shown for structure populations generated with setups H and HDSF.

Interestingly, models from Hi-C data alone (setup H) reproduce SON TSA-seq data and SAF already with high accuracy, while lamina-B1 DamID and LAF show relatively poor performance (**Fig. 4**), which is likely related to the flat ellipsoid shape of the HFF nucleus. Our previous studies on GM12878 cells, with a spherical nucleus, could predict both lamina TSA-seq and laminB1 DamID data with higher accuracy from Hi-C data alone^30^. When Hi-C and Lamina DamID data (setup HD) are combined, predictions of TSA-seq, DamID data, SAF and LAF greatly improve (**Fig. 4**).

Combining SPRITE colocalization clusters and 3D FISH distance distributions with Hi-C and lamina-B1 DamID input information slightly improves correlation scores for TSA-seq and DamID data, even though the total number of spatial restraints from DNA SPRITE and FISH data are an order of magnitude smaller than those from Hi-C and lamina DamID (**Extended Data Figure 2d**). Models HDS and HDSF recapitulate MERFISH imaging data well, recapitulate 3D FISH and SPRITE data, while also showing excellent predictability for TSAseq and DamID data (**Figure 4**, **Extended Data Figure 3**). Overall, the steady improvement of model accuracy with increasing amount of input data highlights the benefits of multi-modal over uni-modal data integration in generating realistic and highly predictive structures.

### Systematic assessment of comprehensive data integration using synthetic data

In order to perform a thorough assessment of multimodal data integration, we regard a structural population as a “ground truth” reference, from which a variety of synthetic data can be simulated (*Methods*) (**Fig. 5a**). Models are then generated from different combinations of synthetic data, to facilitate the comparison of their predictive power on 3D genome architecture. Note that model assessment depends on the structural features being explored, and a ground truth allows a more comprehensive model validation based on a larger number of structural observables that are accessible. For instance, we can now compare how well distributions of distances between all genes are reproduced or how well chromosome shapes, defined by their radius of gyration, match those in the ground truth, which is not feasible with real data. Moreover, we can simulate different input data at variable information content to better assess its influence on model quality.

**Figure 5.**
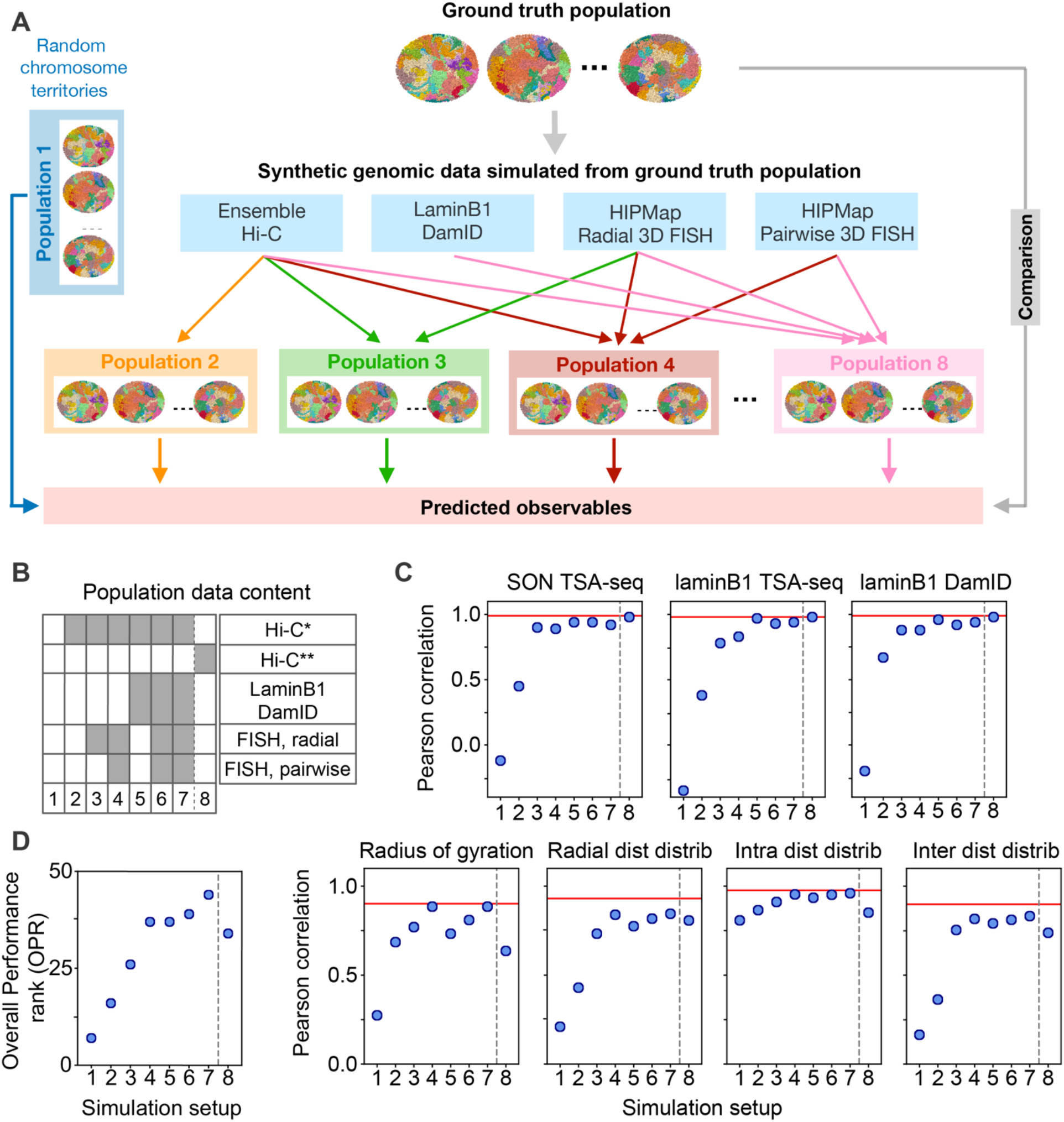
Systematic data integration via synthetic genomic data. (a) Schematic view of the assessment process. Information corresponding to Hi-C, lamina DamID, radial and pairwise FISH data is simulated from a structure population that serves as a reference ground truth. 8 different genome structure populations are calculated from different combinations of synthetic data. Independent structural features are calculated from each population and compared with the ground truth reference to assess the accuracy of the models. (b) Combinations of synthetic data included in 8 different input setups (columns). Dark boxes indicate the presence of a synthetic data type in the input setup. Hi-C* and Hi-C** indicate two differently perturbed Hi-C maps. In Hi-C* only inter-chromosomal contact frequencies are scaled down by a factor of 2. In Hi-C** only intra-chromosomal contact frequencies are scaled down by a factor of 2. More details are provided in the text. (c) Accuracy of models generated from each input setup. Pearson’s correlations between predicted structural features and those in the ground truth reference. Structural features are SON and laminB1 TSA-seq data, laminB1 DamID data, the radius of gyration for chromosomes, distributions of chromatin radial positions, distributions of intra-chromosomal distances, and distributions of inter-chromosomal distances. Baseline predictions from the correct (non-perturbed) Hi-C only simulation are indicated with a red horizontal line. (d) Overall performance ranks (OPR) for all setups. The OPR for setup *s* is calculated as follows: 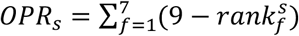, where 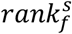 is the rank of setup *s* in assessment of feature 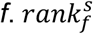 is 1 for the top-ranking setup, and 8 for the poorest performing setup for feature *f*. Therefore, overall performance ranks can range from 56 (top performing for in all feature assessments) to 8 (poorest performance in all feature assessments).

We choose population H (i.e., generated from experimental Hi-C data, see **Fig. 4**) as the *ground truth* structure population, from which we generate the synthetic datasets, including genome-wide contact frequencies (i.e., Hi-C data), contact frequencies between loci and the nuclear envelope (i.e., lamina-B1 DamID data), and a randomly chosen subset of 1000 radial and 1000 pairwise distance distributions (i.e, HIPMap 3D FISH datasets) (Methods) (**Fig. 5a**). These data sets represent idealized data sources, and were combined into seven different input data setups. Models were then generated for all data setups, each containing different combinations of synthetic data (**Fig. 5b**).

We quantitatively assess model accuracy with the following structural properties (**Fig. 5c**): (i) the distribution of radial positions for each chromatin region, (ii) the distributions of pairwise distances between chromatin loci in cis and trans; (iii) the distribution of the radius of gyration for each chromosome; (iv) SON-TSA-seq data, (v) laminB1 TSA-seq data and (vi) laminB1 DamID data. We used the cross-Wasserstein Distance to measure the similarity between two probability distributions (for features i-iii); quantities (iv-vi) are assessed by their Pearson correlation with the corresponding ground truth features (*Methods*). Finally, for each setup an overall performance rank (OPR) is determined as the total sum of ranks for all individual feature assessments (**Fig. 5d**).

Models generated from simulated contact frequencies naturally reproduce with high accuracy the ground truth features. To better substantiate our assessment of data integration performance, we manipulated the simulated Hi-C data by scaling down the inter-chromosomal contact probabilities by a factor of two and used the resulting “perturbed” contact map (labelled Hi-C*) as input for all model populations instead.

Structures generated from perturbed Hi-C* data alone (setup 2) show poor performance with low correlations of ground truth features, except for intra-chromosomal distance distributions (Pearson correlation 0.79) (**Fig. 5c**). We then generated another perturbed Hi-C** data set, in which inter-chromosomal interactions remain untouched, while probabilities of intra-chromosomal interactions are scaled down by a factor of 2 (setup 8). Models generated with this data set predict with good accuracy all ground truth features related to the global nuclear architecture, such as SON TSA-seq, lamina-B1 TSAseq and lamina DamID signals (Pearson correlations > 0.98) as well as radial distributions of chromatin regions with substantially higher accuracy than setup 2 Hi-C* (**Fig. 5c**). In contrast, setup 8 shows slightly higher accuracy than setup 2 for chromosomal properties, such as the radius of gyration. It is noteworthy that intra-chromosomal distance distributions are still well reproduced in comparison to setup 2, which indicates that scaling down intra-chromosomal contacts has a less detrimental effect than inter-chromosomal contacts. These results showcase the surprisingly dramatic loss of information when trans contact probabilities are underestimated in Hi-C data, which generally have very low contact probabilities to begin with. Reducing inter-chromosomal interactions further will lead to the loss of information about the global genome architecture. Reducing relatively high frequent intra-chromosomal contact probabilities has a smaller impact, as sufficient information about intra-chromosomal chromatin interactions is still retained in the data set.

To further assess the relevance of inter-chromosomal interactions, we generated four structure populations from (unperturbed) Hi-C data that include inter-chromosomal contacts only if their contact probability is larger than a given cutoff *θ_inter_*, which is gradually decreased (*Methods*). Interestingly, good predictive models can only be generated when inter-chromosomal contacts with very low probabilities are included (**Fig. 6**). For instance, radial profiles are only reproduced with low residual errors if relatively “rare” contact events are included, i.e., probabilities corresponding to only 2 contact events per 1000 models (**Fig. 6a**). To further substantiate this observation, we also calculate the chromatin compartmentalization score, which measures the spatial segregation between chromatin in the active A compartment from the inactive B compartment^66^ (Methods). The compartmentalization score steadily increases when inter-chromosomal contacts with low contact probabilities are added (**Fig. 6b**). Thus, the large number of low probability inter-chromosomal interactions, which define relatively “rare” contact events per chromatin region, are essential for accurate genome structure modeling and for correct predictions of genome-wide SON TSA-seq, laminB1 TSA-seq and laminB1 DamID data (**Fig. 6c**). Overall, these results further underline the important role of trans interactions in predicting the correct global genome architecture in our models. Hi-C experimental conditions can influence fragment lengths, ligation efficiencies and thus the amount of informative inter-chromosomal proximity information captured by ligations. Hi-C variants, such as MicroC^6^, capture local short range chromatin interactions at higher resolution, while the fraction of long-range and inter-chromosomal interactions is reduced. It is therefore of interest to test if additional orthogonal data sources can compensate for reduced levels of informative inter-chromosomal interactions.

**Fig. 6.**
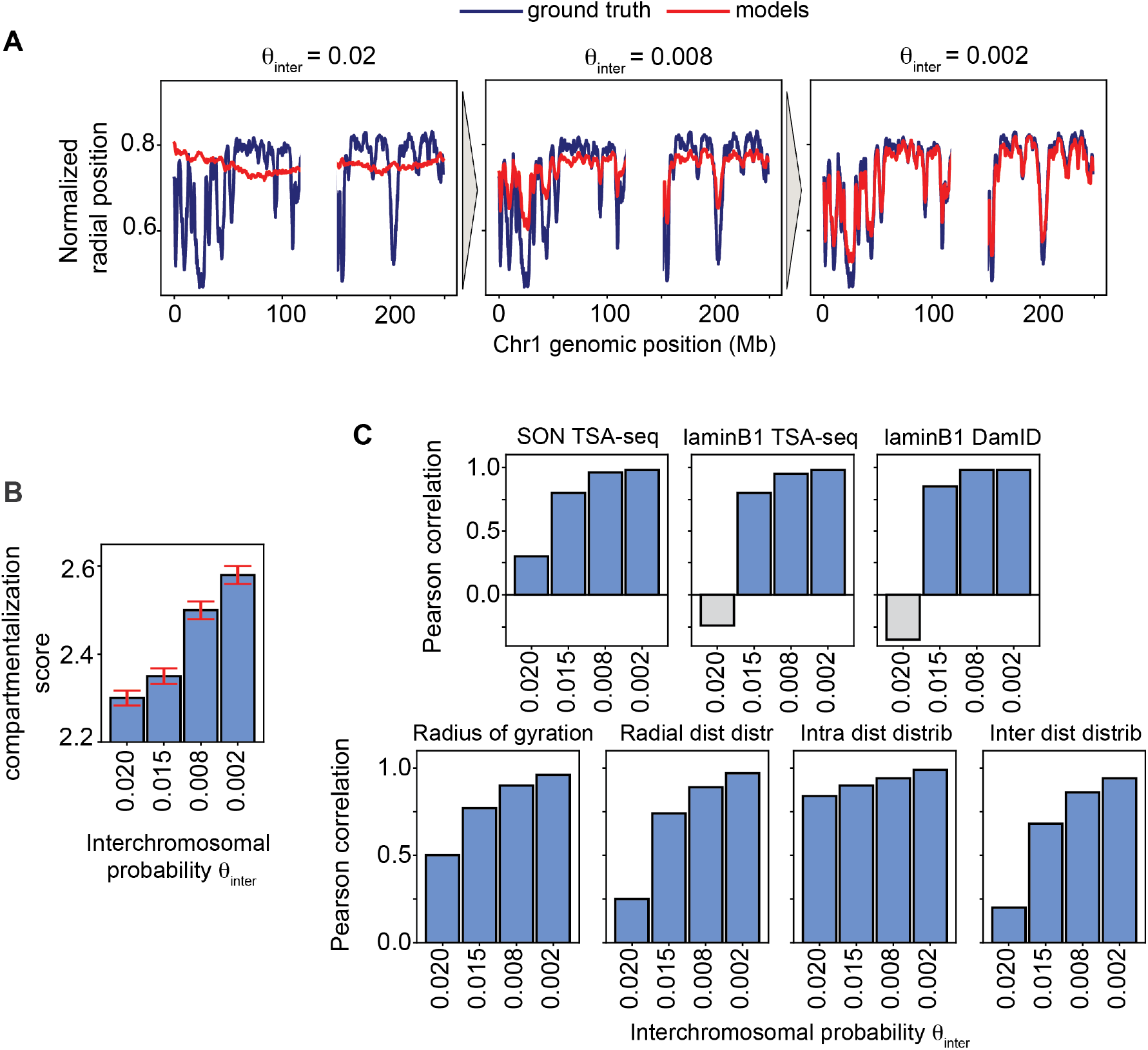
Low probability inter-chromosomal contacts greatly affect model predictivity. We compare the accuracy of a structure population generated with unperturbed Hi-C data as a function of the lowest probability value with which inter-chromosomal contacts are included in the modeling. The probabilities are labeled as *θ_inter_* (*Methods*) (a) The mean radial positions plotted for all chromatin regions in chromosome 1 for structures in the ground truth reference population (dark blue) and structures in populations calculated from three representative setups (red) that include inter-chromosomal contacts with gradually decreasing contact probabilities: *θ_inter_* = 0.02, 0.008, 0.002. Characteristic radial profiles in the ground truth (see also **Fig. 1**) are only reproduced when contacts with a probability smaller than 0.8% are included, and the profiles gradually converge to the ground truth as the probability decreases further. Radial profiles are only correctly reproduced when contacts are included with probabilities of at least 0.2%. From left to right, *θ_intra_* = 0.02,0.008,0.008. (b) The A/B compartmentalization score is plotted for each setup, with error bars representing the standard deviation of the underlying distribution (*Methods*): compartmentalization increases as more low frequency inter-chromosomal contacts are included in the modeling. (c) The Pearson’s correlation value between the ground truth and simulations of the same seven structural observables discussed in **Fig. 5** for *θ_inter_* = 0.020,0.015,0.008,0.002. Grey boxes indicate negative correlation values. Structural quantities experience a substantial correlation increase when low probability contacts are included, which indicates that overall model predictivity increases substantially.

Combining lamina-B1 DamID as well as radial and pairwise distance distributions from 3D FISH experiments with the biased Hi-C* data (setup 7) produces models with high predictive power and similar accuracy for all structural features as models generated with unmodified original Hi-C data (**Fig. 5c**). The overall performance rank increases monotonically with increasing amount of added data (Setups 3-7, **Fig. 5d**). Therefore, orthogonal data modalities appear to compensate for systematic errors affecting one of the data types (here, underrepresentation of inter chromosomal contacts), see also **Extended Data Figure 4**.

The steady improvement in model accuracy with increasing data is not only due to those features being directly restrained by the added data (which is only a small portion of all degrees of freedom), but also due to cooperative effects acting on the entire genome: each newly added data modality makes already included data more informative. This is due to the specific nature of our iterative optimization process, which reduces data ambiguity by selecting the best of a set of alternative restraints assignments, based on the current genome structures at a given iteration (*Methods* and *Supporting Information*). For instance, if newly added information about a gene’s radial position restricts its nuclear locations, it will also make certain non-native chromatin contacts less likely, which in turn will lower the change for that gene to be wrongly selected in non-native Hi-C contact restraint assignments. An analogy is a crossword puzzle, where gradually filling in interconnected words reduces the ambiguity of missing word solutions. Adding a data modality to our modeling process reduces, in a similar way, the ambiguity of restraints assignments of all other data types, thus making these data more informative.

Our simulations show that adding FISH radial distributions for 1000 loci (setup 2 to setup 3) improves prediction accuracy of radial distributions for all genes (not only those being actively restrained), as well as genome-wide SON and lamina-B1 TSA-seq signals, and even inter-chromosomal gene distance distributions, although the radial FISH data do not contain any bivariate information (**Fig. 5c**).

Models generated from Hi-C* and simulated DamID data (setup 5) outperform models from Hi-C* data and FISH radial distributions of 1000 loci (setup 3). However, adding also information for 1000 pairwise FISH distance distributions (setup 4) produces models as accurate as setup 5.

The information equivalence of data sets depends naturally on the amount of data. For instance, using radial distributions of all chromatin loci would render lamina DamID data redundant. We therefore assessed (Hi-C* + radial FISH data) class models that contain increasing numbers of FISH probes. Our results confirm that at a critical number of probes, models from Hi-C* and radial FISH data become more informative than those from Hi-C* and lamina DamID data (setup 5) (**Extended Data Fig. 5**). Of course, these observations are made in an idealized case, and only serve as a conceptual point. The true information content of data depends on systematic errors in the experimental data, such as potential distortions due to cell fixations and other treatments in FISH experiments, as well as the base-pair resolution of the chromatin fiber representation. Also, radial positions (instead of distance to the nuclear lamina) may be an inadequate description for highly irregular nuclear shapes that vary in size. In future, actual microcopy 3D images, instead of positional metadata, shall be used in the modeling process to overcome some of these issues.

## Discussion

We introduced a robust pipeline for multi-modal data integration to determine 3D structures of whole diploid genomes. These structures revealed a wealth of information about the structural organization of genomes over multiple length scales, along with dynamic variabilities of structural features between individual cells. Collectively these features define the nuclear microenvironment of genes on a genome-wide scale, which can directly be linked to their functional potential in gene transcription and subnuclear compartmentalization^43^. Our method therefore provides an ideal analytical tool for comparative genome structure analysis, which could link changes in a gene’s structural organization between different cell types (or during developmental processes) with underlying functional changes. Moreover, the structures generated by our method also predict a host of orthogonal experimental data, including SON TSA-seq data, speckle and lamina association frequencies and trans A/B ratios as determined by DNA-MERFISH experiments. These predictions could serve as first approximations to data otherwise only available through experiments with considerable added effort.

We tested the proficiency of our approach by studying the diploid genome structures of human HFFc6 cells by integrating data from Hi-C, lamin-B1 DamID, 3D HIPMap FISH and SPRITE experiments. The method is generally applicable to any cell type and organisms, with any combination of data available, discussed here. We systematically assessed the accuracy of models generated from different combinations and amount of data types. Model accuracy is steadily improving with increasing amount of data and is maximal when data integration is multimodal, indicating that single data sources might not fully capture all information about a genome’s structural organization. Moreover, orthogonal data sources can compensate for systematic biases and missing information in some data types. For instance, a biased Hi-C data set with artificially reduced chromatin interactions frequencies shows substantially lowered accuracy. However, combining this biased data set with additional information from lamina DamID and 3DFISH experiments recovers structures with almost identical accuracy to those generated by the unbiased Hi-C data. The improvement of performance can partly be explained by cooperative effects. Adding a complementary data type to the input set can reduce ambiguity in other data, thus making already included data more informative.

It is also noticeable that different combinations of orthogonal data sources can produce models with similar levels of high accuracy and thus share similar information content. For instance, the combination of Hi-C with lamina DamID data can produce similarly accurate structures than a combination of data from Hi-C and 3D FISH experiments, given that a critical number of FISH probes is considered. Therefore, the method does not rely on a specific combination of data to produce models with high predictive values.

Interestingly, our work also underlines the essential role of low probability inter-chromosomal interactions for accurate data-driven predictions of genome organizations. The multitude of relatively “rare” contact events are crucial for accurate predictions of radial gene positions and overall chromatin compartmentalization. It is not sufficient to consider only the most significant interactions in the modeling process. However, if data sets are compromised by a lack of sufficient information about trans interactions, additional orthogonal data sources can compensate for reduced level of information.

In future, our approach will be expanded to incorporate also 3D imaging data into the modeling process, which will consider variations in nuclear shapes between individual cells and excluded volumes for some nuclear bodies. We expect that these additions will further improve the quality of models. Due to its modular organization, our software platform is ideally suited for incorporating new volumetric microscopy data

In summary, here we showed that our method provides a useful tool for multimodal data integration to produce genome structure models with high predictability. Our software implementation is publicly available, widely applicable to any cell type, and due to its modular design can be tailored to include new experimental data types.

## Acknowledgements

This work was supported by the National Institutes of Health (grants U54DK107981 and 1UM1HG011593 to F.A), and an NSF CAREER grant (1150287 to F.A.). We thank the laboratories of Profs. Job Dekker (University of Massachusetts Medical School, UMass), Bas Van Steensel (Netherlands Cancer Institute, NKI), Tom Misteli (National Institutes of Health, NIH), and Andrew Belmont (University of Illinois Urbana-Champaign, UIUC) for kindly providing the experimental data (*in situ* Hi-C, lamina DamID, 3D HIPMap FISH, DNA SPRITE and SON TSA-seq, respectively) used for generating and validating our genome models.

## Extended Figures

**Extended Data Figure 1.**
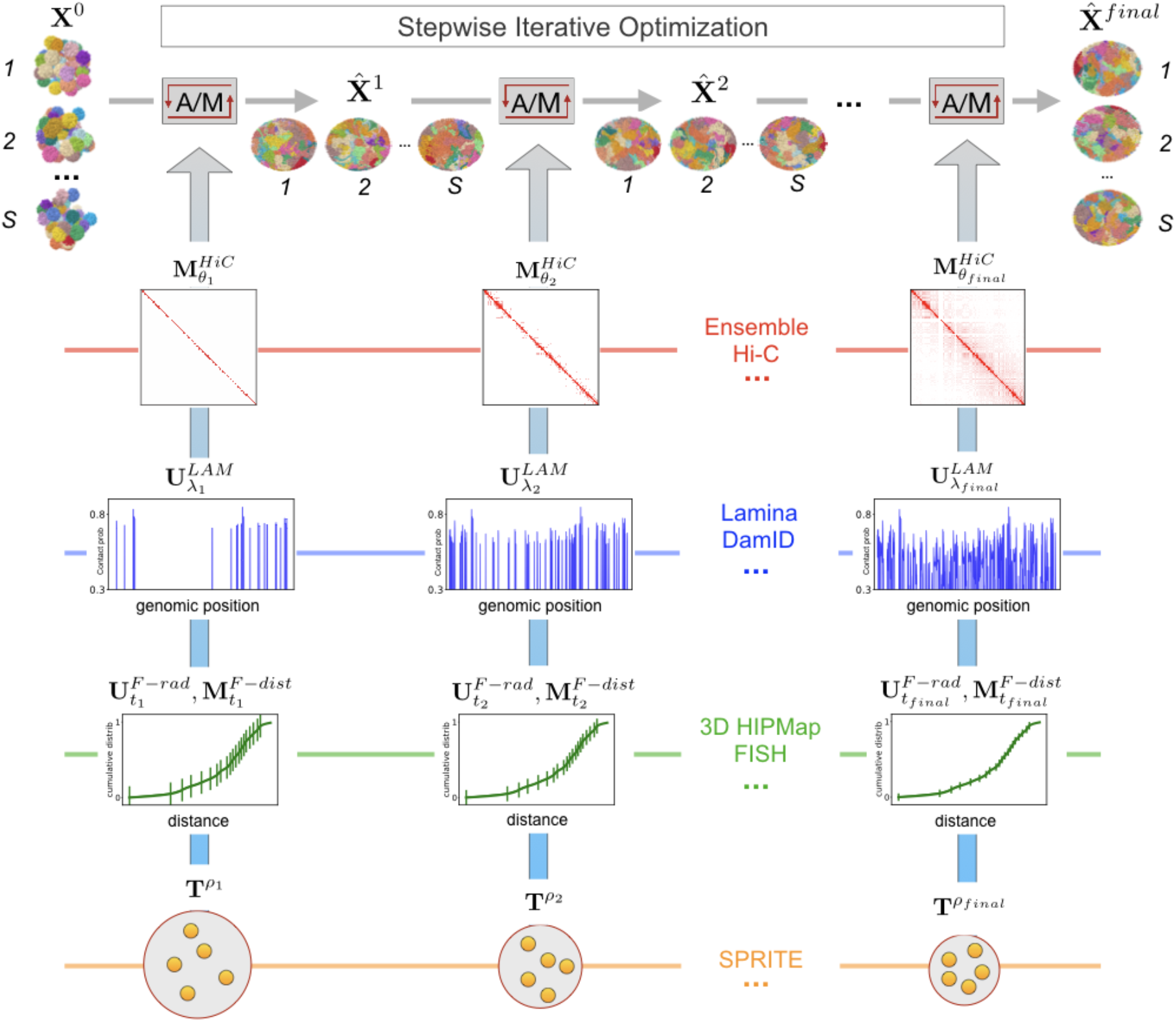
Flowchart of the Stepwise Iterative Optimization pipeline. Ensemble Hi-C, lamina DamID, 3D HIPMap FISH and SPRITE data are used as input to the Stepwise Iterative Optimization protocol which underlies the Integrated Genome Modeling platform. A randomly initialized diploid genome population with chromosome territories ***X***^0^ is first thermally relaxed subjected to envelope and polymer restraints only (not shown). Then, genomic data are gradually added and structures are optimized via a sequence of iterative A/M optimization steps. Optimization hardness is gradually increased by adding batches of data and reducing the tolerance, as visually indicated (see also *Methods*). For example, at the end of i-th A/M step, all contacts with probability larger than ***θ***_*i*_ (i.e., all matrix entries specified by 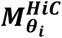), all lamina contacts with probability larger than ***λ***_*i*_ (i.e., all entries 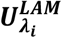), all 3D HIPMap FISH distances with a tolerance equal to ***t***_*i*_ (i.e., 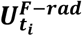 and 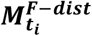) and all SPRITE clusters with volume density ***ρ***_*i*_ (i.e. ***T***^***ρ***_*i*_^) are included (see *Methods*). Multiple sequential A/M iterations may be needed for a given set of optimization thresholds in order to generate an intermediate population 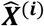 which successfully incorporates all the data restraints that have been added up to that point. At the end of the pipeline, all data up to the final threshold values are included, and, after additional iterations lead to convergence (all data is satisfied), the optimized population 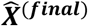 is returned, together with the final violation statistics (see also **Extended Data Figure 2**).

**Extended Data Figure 2.**
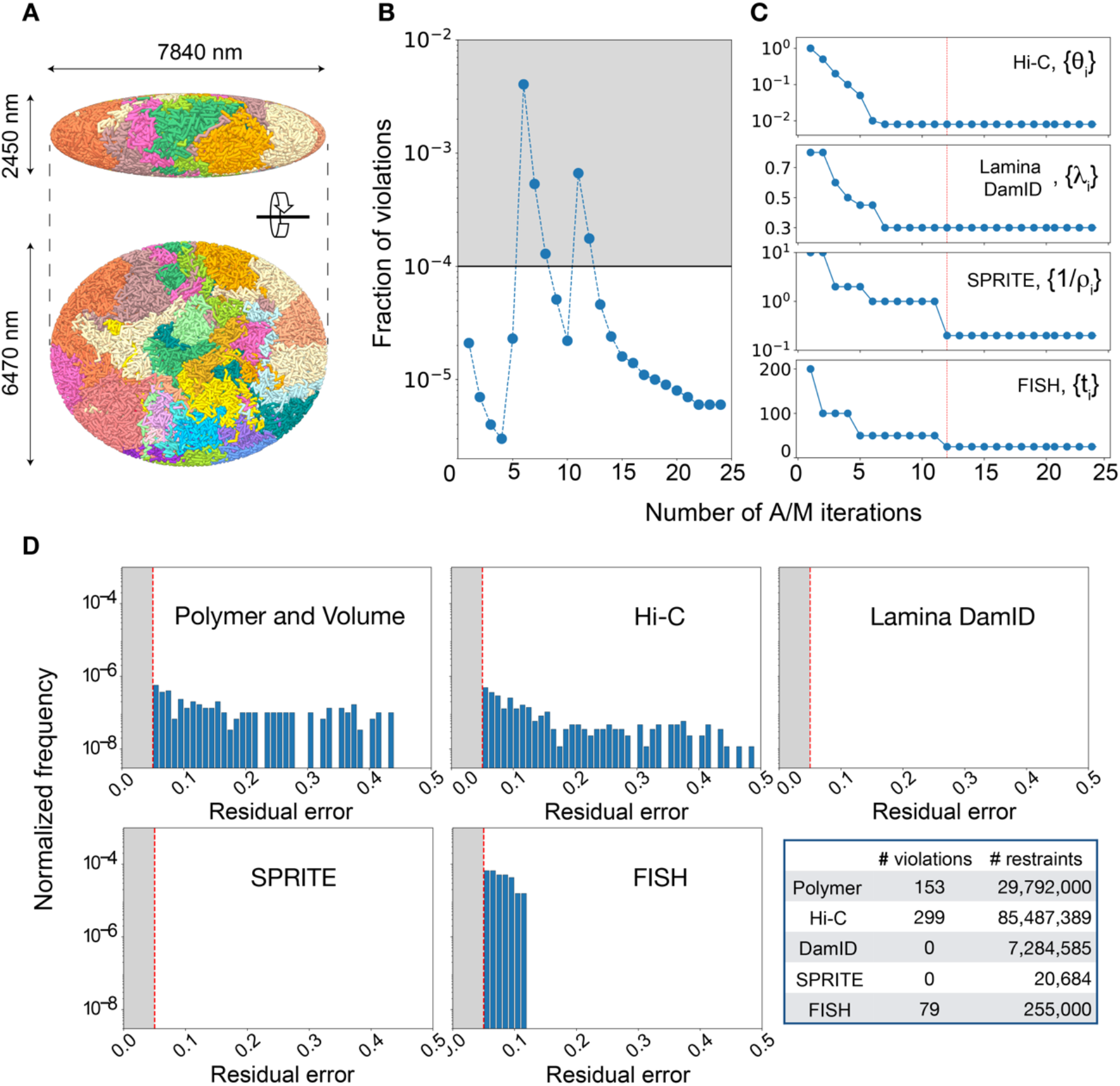
Optimization statistics for HFFc16 all-data genome model. (a) Top and side view of one full genome structure from the optimized HDSF population, with the ellipsoidal nuclear lamina axes annotated (in nm): the same color is used for homologous chromosomes. (b) Fraction of violations plotted as a function of A/M iterations during the HDSF population optimization: jumps in the curve (iterations 6 and 11) indicate the gradual addition of more data batches (i.e. data added at optimization thresholds (Methods)). All data are added by iteration 12, but additional iterations are run to ensure robust convergence with a violation fraction < 10^−5^. (c) Optimization thresholds (*θ_i_,λ_i_,t_i_* and 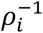), which control the rate and size of data batches being added, shown as a function of the number of A/M iterations: a red vertical line indicates the iteration when all data points are added to the modeling. Final values are non-zero, which reproduces typical experimental setups where finite precision is only available. 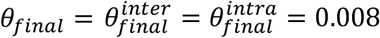 (Hi-C probability), *λ_final_* = 0.3 (lamina DamID probability), *t_final_* = 25*nm* (FISH distance tolerance), *ρ_final_* = 0.005*nm*^−3^ (SPRITE volume density), see also *Methods* and **Extended Data Figure 1**. (d) Final violation statistics broken down into the different restraint categories; each panel shows the normalized histogram of residual errors (*η* > 0.05, see *Supplementary Information*) associated with violations in a given data category. No bars are showing in the SPRITE panel because all applied SPRITE restraints are satisfied, and none is violated. The accompanying table details the number of applied restraints and the number of violations: over 99.999% of polymer restraints, over 99.999% of Hi-C restraints, 99.98% of FISH restraints, and 100% of both SPRITE and lamina DamID restraints are satisfied in the optimized population. The number of FISH and SPRITE restraints is orders of magnitude smaller than polymer, Hi-C and DamID restraints.

**Extended Data Figure 3.**
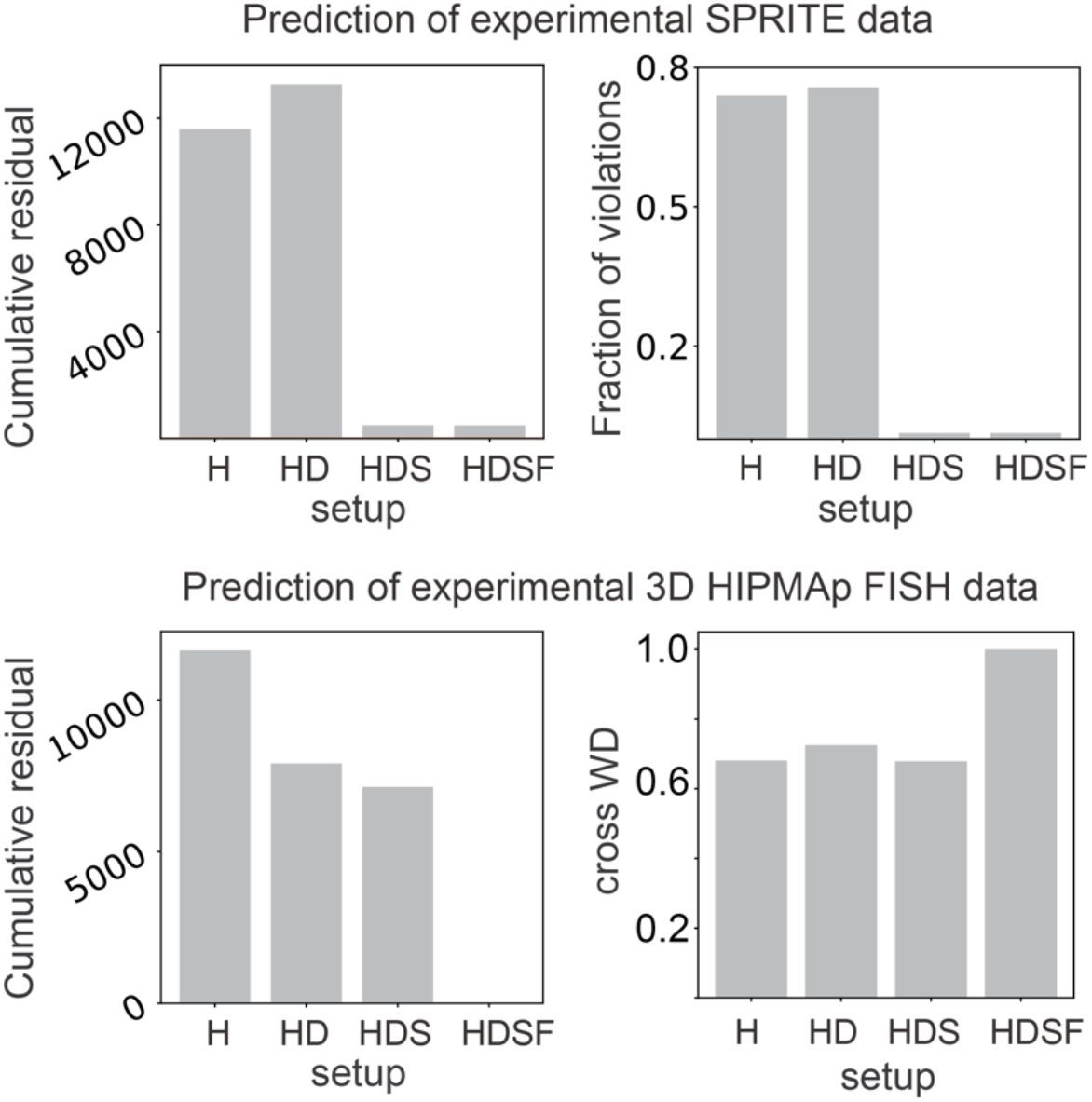
Prediction of experimental SPRITE and FISH data in HFFc6 H, HD, HDS, HDSF populations. (Top panels) SPRITE^11^ cumulative residual (left) and fraction of violated SPRITE restraints (right) for each of the data-driven populations discussed in **Fig. 4**. Lamina DamID restraints tend to stretch the genome towards the lamina, whereas SPRITE restraints squeeze the targeted loci close to one another: an optimal balance is only found when both data modalities are simultaneously integrated, e.g., populations HDS and HDSF. (Bottom) FISH cumulative residual (left) and cross WD score (right). The cumulative residual is defined as the sum of the residual errors *η* for all violations; the cross WD score is the Pearson correlation between two cross WD sets (see *Methods* and *Supporting Information*). FISH distributions^59^ are gradually better predicted with increasing amount of data and most efficiently recapitulated in population HSDF only, as suggested by a cross WD score of 0.999 and the smallest cumulative residual.

**Extended Data Figure 4.**
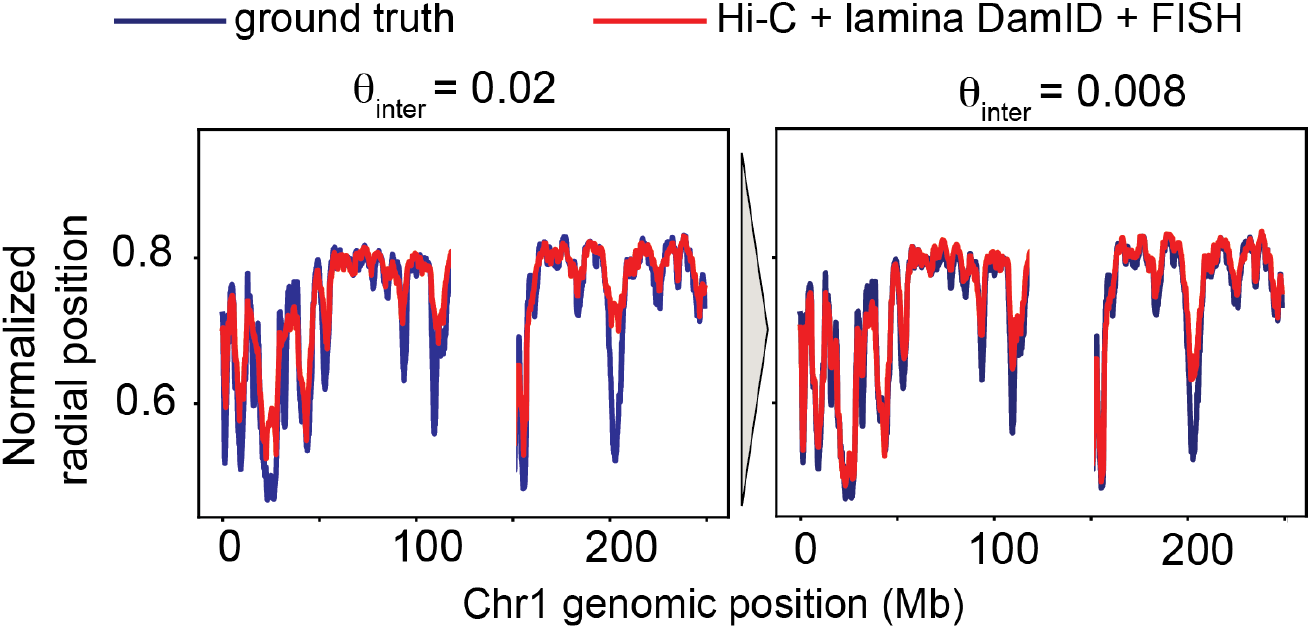
Relevance of low frequency inter-chromosomal contacts. (Unperturbed) Hi-C, lamina DamID and 1000 radial and 1000 pairwise FISH distance distributions extracted from the ground truth (**Fig. 5**) are used to generate a population of structures. The predicted radial profiles for chromosome 1 are compared with the underlying ground truth at different stages of the optimization process. Specifically, lamina DamID and FISH data have been all added up to the final thresholds *λ_final_* and *t_final_*, and low frequency inter chromosomal contacts added up to probability *θ_inter_* = 0.02 (left) and *θ_inter_* = 0.008 (right). Radial profiles are better reproduced in multi-modal Hi-C + lamina DamID + FISH models at *θ_inter_* = 0.02 than they are in Hi-C only models with the same setup (**Fig. 6a**), and then refined by lowering the contact probability *θ_inter_*. This provides alternative evidence that independent data sources can account for missing information; here, inter chromosomal contacts with probability smaller than 0.008. (*θ_intra_* = 0.02,0.008).

**Extended Data Figure 5.**
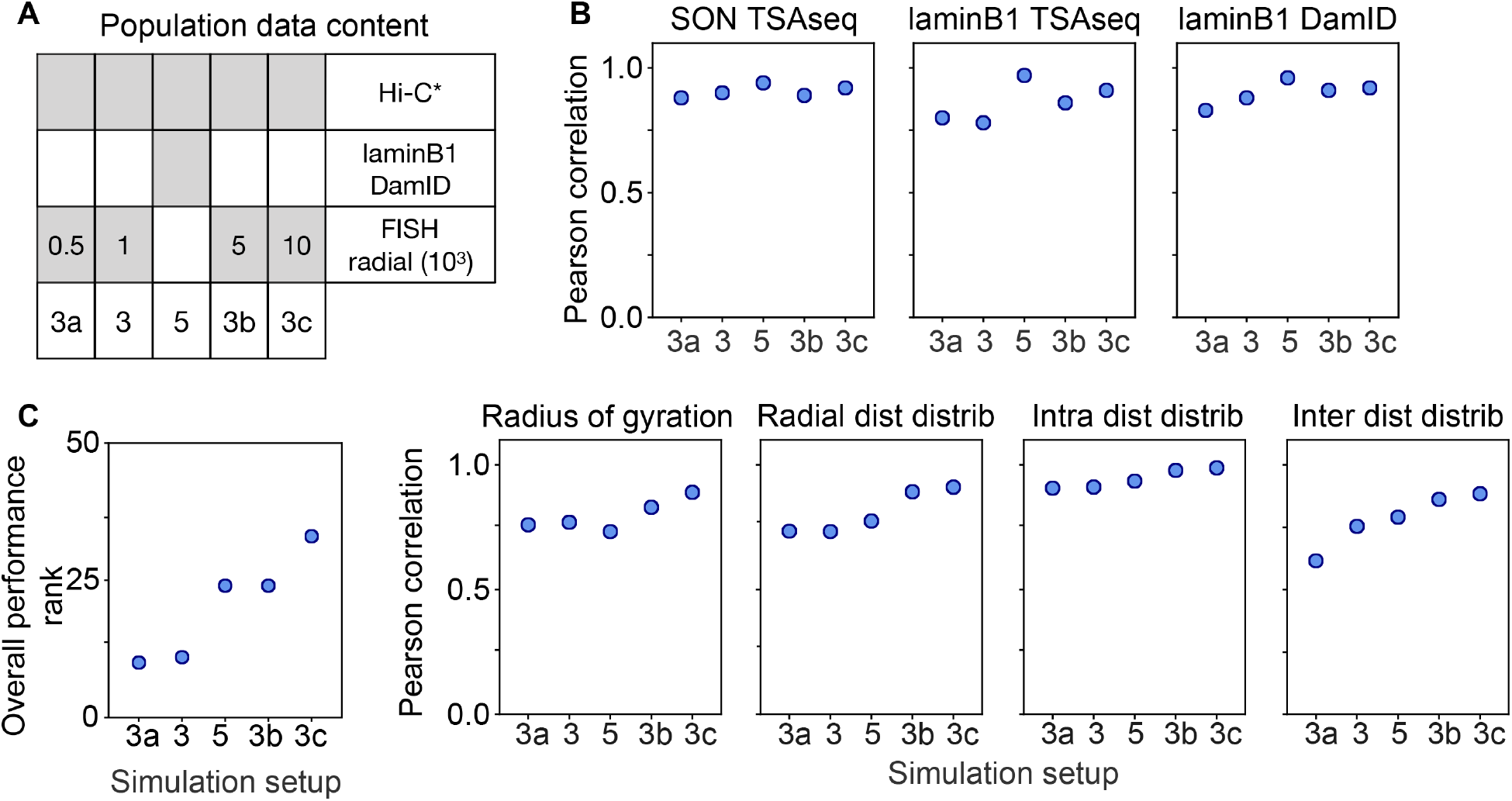
Comparing information content of lamina DamID data against increasingly larger radial distance distribution FISH data sets. Additional Hi-C* and radial FISH only populations (3a, 3b and 3c) are analyzed and compared with previous Hi-C*-radial FISH population 3 and Hi-C*-DamID only population 5 from **Figure 5**. (a) The four populations with FISH data differ in the number of radial distributions used in the input (500, 1,000, 5,000 and 10,000). (b) The seven quantities from **Figure 5c** are predicted for each population and compared with the ground truth. (c) The overall performance rank for these five populations indicates that a sufficiently large sample of radial distance distributions can match and outperform the information provided by lamina DamID data.

## Methods

### 1. Genome representation

A population of S structures at 200kb base pair resolution is a set of *S* diploid genome structures ***X*** = {***X***_1_,…, ***X***_*S*_}; A structure ***X***_*S*_ is a set of 3-dimensional vectors representing the center coordinates of each locus 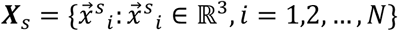, *N* is the number of diploid chromatin regions at a given base pair resolution. In a diploid genome, each autosome chromatin region has two homologous copies. Here, each locus constitutes a 200 kb genome segment, represented by an excluded volume with a sphere radius *r*_0_ = 118*nm*, which guarantees a 40% genome volume occupancy of the nucleus. Overall, the genome is represented by a total of *N* = 29,838 spheres. The nucleus is modeled as a prolate ellipsoid of semi axes (*a, b, c*) = (7,840; 6,470; 2,450) *nm*, see **Extended Data Figure 2a**. The semi axes’ lengths are based on the estimates from Seaman *et al*.^67^.

In the following, we will use uppercase (lowercase) letters for haploid (diploid) indexing. Specifically, (*i, i*′) indicate the two diploid copies associated to haploid locus *I*.

### 2. Data sources and probabilistic formulation of the structure population optimization

Our goal is to generate a population of *S* genome structures ***X*** that are statistically compatible with all available data from different experimental sources. These data sources can be conveniently categorized into classes depending on the number of genomic loci being involved. For instance, any data type that depends on the coordinates of only a single locus will be *univariate*: a locus’ radial distance, normal distance to the nuclear lamina or average distance to speckle clusters, etc., are examples of univariate data. *Bivariate* data involve two genomic loci, for instance distances between pairs of loci, while *multivariate* data defines a relationship between more than 2 loci, for instance knowledge about multivalent interactions co-occurring between several loci.

The four data sources that we discuss in this work are in-situ Hi-C^63^ and lamina DamID^64^, high-throughput HIPMap fluorescence in situ hybridization (3D FISH)^59^ and Split-pool Recognition of Interactions by Tag Extension (SPRITE)(Quinodoz et al., 2018): all data can be interpreted in terms of probability distributions of distances. Let us assume all the accessible distance values (with respect to any reference point) are discretized into *Q* bins and let *d_q_* denote the distance associated with the *q*-th bin.

- **Univariate data** informs a single locus, such as information derived from radial 3D FISH or lamin-B1 DamID data. We express radial 3D FISH data with the tensor 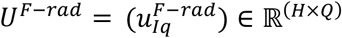, with *H* as the total number of genomic regions and *Q* as the total number of associated distance bins. 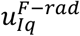 is derived from the HIPMap data and is the probability that the radial distance of locus *I* is equal to the distance *d_q_* in the population. For FISH data, the distances *d_q_*, associated to bins *q*, are equally distributed and span the nuclear dimension. Lamina DamID data is expressed in a similar way by the tensor 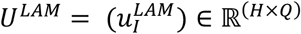 (here, *Q* = 1). Here, 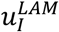 is derived from lamina DamID data and defines the probability that chromatin region *I* is in contact with the lamina at the nuclear envelope (NE), i.e. the distance between *I* and the NE *d*_*q*=1_ < *l^contact^* with *l^contact^* as a lamina contact distance threshold.^38^.
- **Bivariate data** inform pairs of loci, for instance pairwise distance distributions from 3D HIPMap FISH or contact probabilities between pairs of loci from Hi-C data. We express 3D FISH pairwise distance data by the tensor 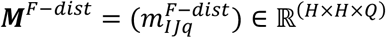, where 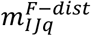 is the probability that the distance between loci *I* and *J* is equal to *d_q_* in the population. We express Hi-C data by the tensor 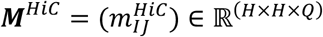 (here, *Q* = 1), where 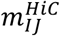 is derived from bulk Hi-C data and describes the contact probability between loci *I* and *J*, thus the probability that the distance between *I* and *J* is *d*_*q*=1_ < *l^C^*, with *l^C^* as a threshold distance defining a contact between *I* and *J*.
- **Multivariate data** inform groups of loci. For instance, SPRITE data provide information that a number of loci form a spatial cluster in a single genome structure. We express the SPRITE data by a collection of tensors 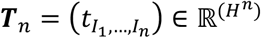, where *n* is the number of loci in an individual SPRITE cluster and *t*_*I*_1_,…,*I_n_*_ is derived from SPRITE data and is the probability of loci *I*_1_,…, *I_n_* to be co-localized in a single structure of the population; *I_g_* is the index of the *g^th^* locus in a SPRITE cluster.

For simplicity, we use ***U, M, T*** to collectively describe all uni-bi- and multivariate data types. With known ***U, M, T*** we calculate the population of structures 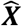 such that the likelihood *P*(***X***) is maximized.

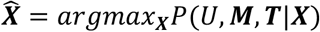

Most experiments, such as Hi-C and Lamina DamID, provide data that are averaged over a large population of cells, and so they cannot reveal which contacts co-exist in which structure. Moreover, all experiments that produce unphased data cannot discriminate between chromosome copies. To represent the missing information at single cell and diploid level, we introduce indicator tensors ***V, W, R*** as latent variables that augment missing information in ***U, M, T***, respectively.

The univariate latent variable 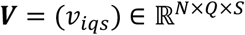 indicates whether the *i*-th diploid locus has a radial distance equal to *d_q_* in structure *s* (*v_iqs_* = 1) or does not (*v_iqs_* = 0). The bivariate latent indicator tensor 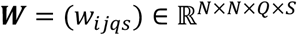 indicates whether the pairwise distance between *i*-th and *j*-th diploid loci is equal to *d_q_* in structure *s* (*w_ijqs_* = 1) or does not (*w_ijqs_* = 0). The multivariate latent indicator tensor 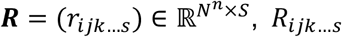 indicates whether the loci *i,j,k*,… are colocalized in structure *s* (*r_ijk…s_* = 1) or not (*r_ijk,…,s_* = 0).

The statistical dependence among all the variables introduced is ***X→W→M,X→V→U,X→ R →T***, since ***X*** is the population consistent with ***V,W,R***, which in turn are detailed expansions of ***U,M,T*** at a diploid and single-structure representation of the data.

### 3. Probability optimization

We formulate the genome structure optimization problem as a maximization of the likelihood that the data sources are expressed in a population.

#### The A/M step

A population that is consistent with all data modalities is identified by searching in the space of genome populations for the optimum 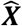, which maximizes the probability *P* that the experimental data are expressed^45,62^.

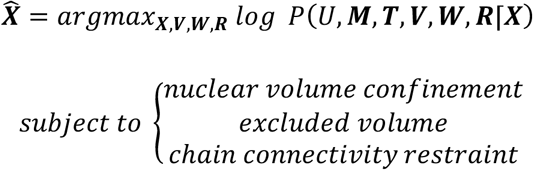

In addition to the genomic data, additional constitutive restraints are applied to reproduce the nuclear volume confinement, chromatin excluded volume and polymer chain connectivity, which guarantees the structural integrity of the chromosomal chains.

The log likelihood can be expanded as

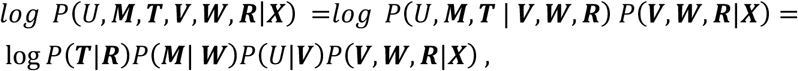

Since there is no closed form of solution to the large-scale optimization problem, we have developed a variant of the Expectation-maximization (EM) method to iteratively optimize local approximations of this log likelihood function^38,45,68^. Each iteration consists of two steps:

- Assignment step (*A-step*): Given the current model ***X***^(*t*)^ at the iteration step *t*, estimate the latent variables ***R***^(*t*+1)^, ***W***^(*t*+1)^ and ***V***^(*t*+1)^ by maximizing the log-likelihood over all possible values of ***V,W,R***:

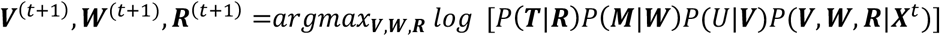 The updated latent variables ***V***^(*t*+1)^, ***W***^(*t*+1)^, ***R***^(*t*+1)^ are optimized given the current structure population at step *t* and describe the optimal allocation of data across the individual structures, and which data instances (e.g. which contacts or which lamina contacts) cooccur within the same structure, based on the structures of the current population at step *t*. We use an efficient heuristic strategy to estimate all latent variables by using information from the structure population generated in the previous *M*-step (*Supporting Information*).
- Modeling step (*M-step*): Given the current estimated latent variables ***R***^(*t*+1)^, ***W***^(*t*+1)^ and ***V***^(*t*+1)^, find the model ***X***^(*t*+1)^ that maximizes the log-likelihood function.

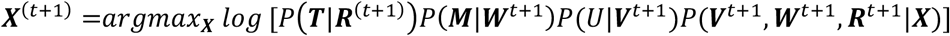
- The effect of each latent variable entry can be modeled as an appropriate spatial energy term (i.e., a spatial restraint) involving one or more genome loci in a given structure (*Supporting Information*). Then, independent structure optimization is performed using a combination of simulated annealing MD^69^ and conjugate gradient^70^ calculations, which is implemented using the open source software LAMMPS^71^ in an efficient parallel platform.

A/M iterations are repeated until convergence is reached. At each A/M iteration, all data allocations are re-evaluated using the current structure population and restraints are re-distributed across the structures. Optimizations are initiated with random chromosome configurations ***X***^0^: chromatin regions are randomly placed in a bounding sphere proportional to its chromosome territory size and randomly placed within the nucleus followed by an M-step to eliminate steric clashes in the structures.

#### Stepwise optimization strategy

We use a stepwise optimization strategy to gradually increase the optimization hardness (**Extended Data Figure 1**). In the initial step, we first calculate a structure population ***X***^*i*^ that integrates only data with the highest probabilities (for Hi-C and and DamID data) and large distance tolerances (for SPRITE and FISH data) (*Supplementary Information*). At each step, we add further data batches with gradually lower probabilities (for Hi-C and lamina DamID), and decreasing tolerances when applicable (for SPRITE and FISH data), and perform several rounds of iterative A/M optimizations until convergence for all data is reached (i.e., all data is reproduced in the models) (see **Extended Data Figure 2b,c**). How the data is added to the optimization at each step and to which accuracy is controlled by a sequence of nonzero threshold values, and each data type is associated with its own sequence.

- *θ*_1_≤⋯≤*θ_final_* indicates the list of Hi-C probability values, such that the i-th step incorporates those contacts that are more probable than *θ_i_*, 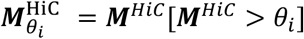.
- *λ_i_* ≤ ⋯ ≤ *λ_final_* is the list of DamID contact probability values such that the *i*-th step incorporates those contacts that are more probable than *λ_i_*, 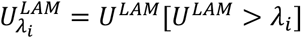.
- *t*_1_≤⋯≤*t_final_* indicates the list of FISH distance thresholds, such that the *i*-th step in the optimization enforces distance values with a tolerance *t_i_*. All FISH distances are incorporated from the first optimization steps on, but their tolerances is gradually reduced with the number of optimization steps.
- *ρ*_1_≤⋯≤*ρ_final_* indicates the SPRITE thresholds, i.e., such that the i-th step enforces clusters with a volume density *ρ_i_*. The volume density is related to the cluster radius, as detailed in the (*Supporting Information*). All SPRITE clusters are incorporated from the beginning of the optimization, while their effective co-location density is increased with numbers of optimization steps.

We use a non-zero final bound for each data type (i.e., *θ_final_,λ_final_,t_final_,ρ_final_* > 0) to reduce the chances of including experimental noise in the calculations (i.e. data errors are expected to have very low probabilities). Multiple A/M iterations are typically necessary at a given optimization step, which is defined by a given combination of threshold values (compare with **Extended Data Figure 2b,c**). Only if optimization in a given step is fully converged the optimization will proceed to the next step. Different data sources are integrated in simultaneously.

The Integrated Genome structure Modeling (IGM) software, we introduce here, automatically performs the sequence of A/M iterations until full convergence is reached and a genome structure populations is calculated that recapitulates all the input data (at a given tolerance) (**Extended Data Figure 1**).

#### Convergence

The optimization progress is monitored by tracking the agreement between model and target distances. As detailed in the *Supporting Information*, each energy term introduced in the M-step to model the effect of genomic data is associated with a residual error *η* which monitors whether the corresponding target distance is satisfied or not: *η* > 0.05 indicates a discrepancy between target and model distances larger than 5%, and is considered a violation. A round of A/M iterations (for a given combination of threshold values) is successful when the cumulative fraction of all violations (from all data types) is smaller than 0.01%. Only then the optimization moves to the next step, and optimization thresholds are lowered and more data are added. **Extended Data Figure 2d** shows the histogram of residual errors in population HDSF for the different data categories used as input (polymer and volume, Hi-C, lamina DamID, SPRITE and FISH).

#### The IGM software

The IGM (Integrated Genome Modeling) requires one input file for each data type and a configuration file, which lists all parameters controlling the pipeline, including nuclear shape, genome segmentation/basepair resolution, nuclear radius, semi axes, MD time step. The software automatically performs a preliminary statistical analysis of genome structures, including a report of the model quality using the correlation between prediction and experiments, and radial features such as the radial positions of individual chromatin domains in the nucleus.

We refer the interested reader to the documentation for implementation details. Here, we would like to discuss the design guidelines that were cornerstones to the development: flexibility, modularity and user-friendliness.

As for flexibility, the software is able to handle different types of genomes confined to either spherical or ellipsoidal nuclei and can use any combination of ensemble Hi-C, laminB1 DamID, 3D FISH and SPRITE data points as an input. Due to IGM’s modularity, the different parts of the code communicate in such a way that any data type can be added with minimal changes, as long as the data can be cast into an energy term, thus allowing for any data customization users may require. Parallel computing can be deployed on different schedulers in a straightforward manner. Simulation and optimization setups can be adjusted by editing a text file, which lists all the configuration parameters.

A python wrapper is available interfacing the different building blocks and keeping tracks of the optimization status.

The optimization progress is monitored by a logger file that prints all the details, from current iteration violation score to the specific values of thresholds associated with it. The complete package (and its documentation) is available at www.github.com/alberlab/igm.

### 4. Simulating structural observables from a population of genome structures

The same notation and variables are used here as in the description above and in the *Supporting Information*. ***x***_*is*_ denotes the 3D coordinates of locus *i* in structure *s*.

#### Ensemble Hi-C

The Hi-C indicator tensor 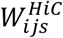 is computed as

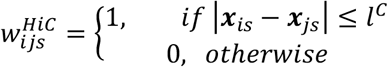

with *l^C^* is the contact distance

The simulated 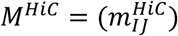 matrix is computed as follows:

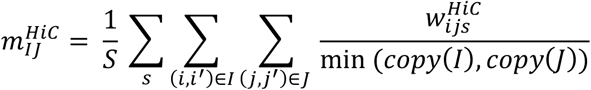

where *copy*(*K*) indicates the number of homologues associated with locus *K*, so that min(*copy*(*I*), *copy*(*J*)) = 1 (if either one of the loci is from a sex chromosome) or 2 otherwise.

#### Lamina DamID

The lamina DamID indicator tensor 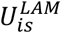 is computed:

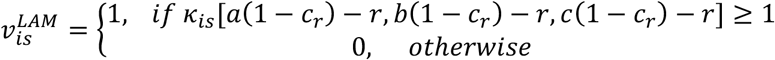

Where *κ* and *c_r_* are appropriate scalars (*Supporting Information*). The simulated 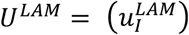 matrix is then computed as: 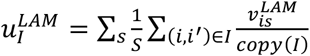

#### Radial and pairwise distance distributions (3D HIPMap)

We extract (radial or pairwise) ordered distance distributions from the *S* structures in the population for the loci of interest:

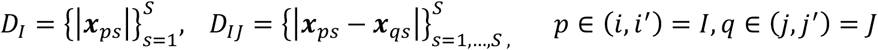

Depending on the number of homologues copies for autosome or sex chromosomes, we end up with *S* or 2*S* radial distances and 2*S* or 4*S* distance values. Those are the target distances we will enforce in a model population of structures, upon filtering and interpolation, and will populate either the *U^F−rad^* or ***M***^*F−dist*^ tensors (see *Supplementary Information*).

#### SPRITE colocalization clusters

For a given SPRITE cluster {*I*_1_,…,*I_n_*}, we follow the Assignment procedure in *Supporting Information*; we compute the SPRITE violation score for all structures: if a structure has no violations, then the cluster is present in that structure, and *t_I_1_,…,I_n__* = 1; If no structure has zero violations, the cluster is not present in the population, i.e.: *t_I_1_,…,I_n__* = 0.

A more detailed description of the following structural features is provided in ref.^30^.

#### Distance of a locus to the nuclear center and to the lamina

The normalized radial distance of a locus *i* of coordinates (*x_is_, y_is_, z_is_*) to the nuclear center of an ellipsoidal nucleus (in population structure *s*) is computed as:

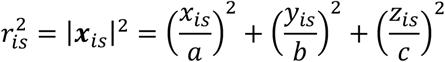

i.e., locus coordinates are scaled by the corresponding semi-axes. |***x***_*is*_| = 0 (1) indicates that the region is located at the geometric center (nuclear lamina).

The normal distance to an ellipsoidal surface cannot be computed exactly, so we use the radial approximation for the distance to the lamina (NE):

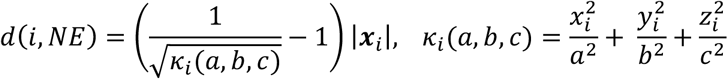

#### Radius of gyration

radius of gyration of a chromatin segment comprising *C* loci *C* = (*i*_1_, *i*_2_,…, *i_C_*) in genome structure *s* is computed as:

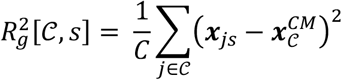

where ***x***_*js*_ are the coordinates of the *j*-th locus in the segment, and 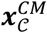 is the segment center of mass in structure *s*. The chromosomal radius of gyration is easily computed by replacing a chromatin segment with a whole chromosome.

#### Compartmentalization score

For HFFc6 cell type, each locus is assigned to either A or B compartments using the ensemble Hi-C and the procedure in following^8^. For each structure, the compartmentalization score is computed as defined in ^66^:

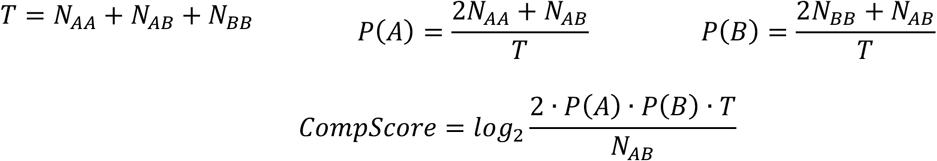

where *N_AA_, N_AB_* and *N_BB_* are the number of A-A, A-B and B-B contacts in the structure respectively. The A/B assignment for HFFc6 structures was downloaded from the 4DN portal^63^ under identifier 4DNFINQZ5JHV.

#### Average radial position

The mean radial position of a locus *I* in an autosome is 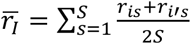, with *i, i*′ as the two homologues copies. *S* is the total number of structures in the population^30^.

#### Chromatin decompaction

The local compaction of the chromatin fiber at the location of a given locus is estimated by the radius of gyration for a 1 Mb regions centered at the locus (i.e. comprising +500kb up- and 500 kb downstream of the given locus). To estimate the RG values along an entire chromosome we use a sliding window approach over all chromatin regions in a chromosome, as described in ref.^30^.

#### Cell-to-cell variability of structural features^30^

Cell-to-cell variability *δ* of any structural feature for a chromatin region, *i*, in chromosome *c*, is calculated as:

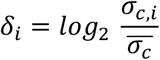

where *σ_c,i_* is the standard deviation of the feature value of region *i* across the population and 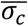 is the mean standard deviation of the feature value calculated from all regions within the same chromosome, *c*. Positive *δ_i_* values (*δ_i_* > 0) result from high cell-to-cell variability of the feature (e.g. radial position); whereas negative values (*δ_i_* < 0) indicate low variability.

#### Inter-chromosomal interaction probability

For each chromatin region *I*, its inter-chromosomal interaction probability (iCP) is calculated as:

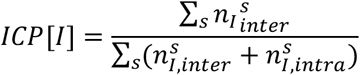

across the full population, where 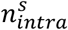 and 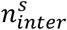 are the number of cis and trans contacts in structure *s*.

#### Interior chromatin localization

For a given 200-kb region, the interior localization frequency (ILF) is calculated as:

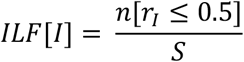

where *n*[*r_I_* ≤ 0.5] is the number of structures where either copy of the region *I* has a radial position lower than 0.5, e.g. is in the nuclear interior.

#### SON TSA-seq

We follow a procedure described in^30^. We first identify chromatin expected to have high speckle association: we select 5% chromatin regions with the lowest average radial positions and generate chromatin interaction network (CINs)^72^ for the selected group of chromatin regions in each structure of the population. A CIN is calculated for the selected chromatin in each model as follows: Each vertex represents a 200-kb chromatin region. An edge between two vertices *i, j* is drawn if the corresponding chromatin regions are in physical contact in the model, if the spatial distance *d_ij_* ≤ 4*r*_0_. Approximate speckle locations are then identified as the geometric center of the resulting spatial partitions identified by Markov clustering^73^ of the CINs.

To predict TSA-seq signals from our models, we use the following equation:

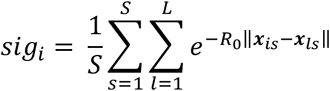

where *S* is the number of models, *L* is the number of approximate speckle locations in structure *s*, |***x***_*is*_ – ***x***_*ls*_| is the distance between the region *i* and the predicted nuclear body location *l* (in structure *s*), and *R*_0_ = 4 is the estimated decay constant in the TSA-seq experiment^61^. The normalized TSA-seq signal for region *i* then becomes:

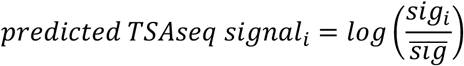

where 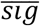 is the mean signal calculated from all regions in the genome.The predicted signal is averaged over copies for regions that have more than one copy in the genome.

#### LaminB1 TSA-seq

We follow the procedure described in^30^. For lamin locations we first identify regions with the highest 15% radial positions in each structure, determine spatial partitions of these regions and use centers of these spatial partitions as approximate locations of lamina associated domains. Lamina-TSA-seq signal is then calculated from these center locations using the decay function described in the SON-TSA-seq section.

#### Speckle (SAF) and Lamina (LAF) Association Frequency^30^

For a given 200-kb chromatin region *I*, the speckle association frequency (SAF) is calculated as:

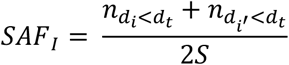

where *S* is the number of structures in the population; *n_d_i_<d_t__* and 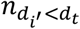 are the number of structures, in which region *i* and its homologous copy *i*′ have a distance to a predicted speckle smaller than the association threshold, *d_t_* (if the chromatin region is from a sex chromosome, there is only one copy and *i′* = *i*). The *d_t_* is set to 1000 nm. Distances to the speckles are computed using the predicted speckle partitions via Markov Clustering which is explained in the section above.

For a given 200-kb chromatin region *I*, the lamina association frequency (LAF) is calculated as:

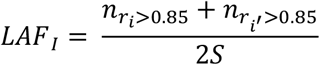

where *S* is the number of structures in the population; *n_r_i_>0.85_* and 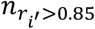 are the number of structures, in which region *i* and its homologous copy *i′* have a radial position larger than 0.85 (if the chromatin region is from a sex chromosome, there is only one copy and *i′* = *i*). Both for SAF and LAF, we try different distance thresholds, and the selected thresholds resulted in the best correlations with experimental data. The following experimental threshold distances were used for comparison with the experimental data from Su et al.^17^: SAF; 500 nm, LAF; 750 nm.

#### Median trans A/B ratio^17,30^

For each chromatin region *i*, we define the trans neighborhood {*j*} if the center-to-center distances of other regions from other chromosomes to *i* are smaller than 500 nm, which can be expressed as a set; 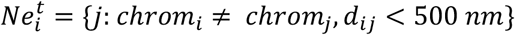. Trans A/B ratio is then calculated as:

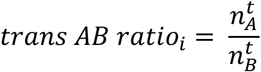

where 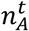 and 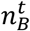 are the number of trans A and B regions in the set *Ne_i_* for haploid region *i*. The median of the trans A/B ratios for a region is then calculated from all the trans A/B ratios of the homologous copies of the region observed in all the structures of the population. The values are then rescaled to have values between 0 – 1.

### 5. Data analysis

#### Correlation

Unless otherwise specified, Pearson correlation is employed in order to compare a given quantity across different populations. All the correlation values given in the text are associated with a p-value < 10^−4^.

#### Cross Wasserstein Distance

Let *Q* and *P* denote the cumulative probability distributions (CDFs) of distributions *q* and *p* of variable *y*, then the Wasserstein Distance^74^

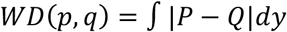

Is customarily used to estimate the amount of work required to transform one distribution into the other, “work” measured as the amount of distribution weight to be moved, multiplied by the distance it has to be moved. We employ the plain WD distance to compare two distributions within the same population.

When comparing probability distributions between two different genome populations or between one population and a set of experimental data, we use the notion of cross (“all versus all”) Wasserstein Distance: we compute the set of all WD for applicable distribution pairs within the same populations (cross WD) and then compute a simple correlation between the two sets (*score*). Let us assume we would like to compare the set of distance distributions of *n* pairs *C* = {(*i*_1_,*j*_1_),⋯,(*i_n_,j_n_*)} between population 1 and population 2 (either one could be an experimental distribution): then we will compute

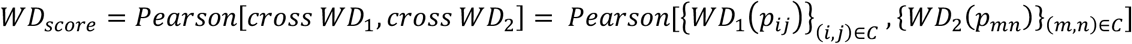

which is the correlation between two sets of *n*(*n* – 1)/2 WD values. Please note that for a given haploid pair *I* – *J* the four diploid pair distributions are concatenated, *p_IJ_* = *p_ij_* ∪ *p_ij_*, ∪ *p_i′j_* ∪ *p_i′j′_*. Cross WD we use to compare distance distributions in **Fig. 2e**, to compare radial, cis and trans pairwise distance distributions, and chromosomal radius of gyration in **Figs. 5c, 6c and Extended Data Figure 4b**.

#### Data and Structure visualization

3D genome models are visualized by using Chimera^75^. Data post-processing was performed using the matplotlib^76^ and Scikit-learn^77^ Python packages.

