## Supplementary Information for "Integrative Genome Modeling Platform reveals essentiality of rare contact events in 3D genome organizations"

* To whom correspondence should be addressed.

### Table of contents:

1. A/M optimization: Assignment Step
    - Univariate data
    - Bivariate data
    - Multivariate data
2. A/M optimization: Modeling Step
    - Univariate data
    - Bivariate data
    - Multivariate data
    - Polymer chain
    - Volume confinement
3. Synthetic data simulations
4. Iterative contact refinement step
5. HFFc6 experimental data pre-processing:
    - Ensemble Hi-C
    - Lamina DamID
    - DNA SPRITE
    - 3D HIPMap FISH
    - SON TSA-seq

#### 1. A/M optimization: Assignment Step

Starting from a population of structures $\boldsymbol{X}^{(t)}$, the latent variables $\boldsymbol{V}^{(t+1)}, \boldsymbol{W}^{(t+1)}$ and $\boldsymbol{R}^{(t+1)}$ for next iteration are fully determined in the Assignment Step (*Methods*). We will briefly detail the heuristics guiding the process.

For the sake of generality, we will assume each haploid locus $I$ comes with two diploid copies $(i, i')$, but the discussion can easily be adjusted for non-autosome single copy loci.

We will use the following notation. $\boldsymbol{x}_{is} = (x_{is}, y_{is}, z_{is}) \in \mathbb{R}^3$ is the coordinate of locus $i$ in structure $s$ of the population, and $\boldsymbol{x}_{ijs} = \boldsymbol{x}_{is} - \boldsymbol{x}_{js}$. We use $| \dots |$ to denote the norm of a vector. $\{x_1^I, \dots, x_S^I\}$ is the ordered discrete radial distance distribution for locus $I$ (from experiment), and $\{y_1^{IJ}, \dots, y_S^{IJ}\}$ is the ordered discrete pairwise distance distribution for locus pair $I$ and $J$ (from experiment). Distances from experiments we will refer to as target distances, since those we would like our models to reproduce and we assume those distances are already pre-processed and mapped onto a set of $S$ distinct values, $S$ being number of structures in the population (see also *HFFc6 experimental data pre-processing*). $r_0$ indicates the locus radius.

Let us assume all the accessible distance values (with respect to any reference point) are discretized into $Q$ bins and let $d_q$ denote the distance associated with the $q$-th bin (*Methods*). Also, let $\sigma[k: K]$ indicate the ranking function that returns the rank of the element $k \in K$ when the t-uple $K$ is sorted in ascending order.

**Univariate data**, locus $I$.

<u>3D radial FISH</u>: By using the information in the data $U^{F-rad} \in \mathbb{R}^{(H \times Q)}$ and the population $\boldsymbol{X}^{(t)}$, we determine the latent variable $V^{F-rad} \in \mathbb{R}^{(N \times Q \times S)}$. The target distances for locus $I$ are given in the list $\{x_1^I, \dots, x_S^I\}$.

We compute the $2S$ radial distances $D_i = \{|\boldsymbol{x}_{is}|\}^S{}_{s=1, i \in I}$ from population $\boldsymbol{X}^{(t)}$, and sort them in ascending order; we obtain a sequence of ordered (diploid, structure) indexes $\{(i_1, s_1), \cdots, (i_{2S}, s_{2S})\}$, such that $|\boldsymbol{x}_{i_n s_n}| \leq |\boldsymbol{x}_{i_m s_m}|, \forall\, n \leq m$. We subsample such list down to $S$ elements (using either a stride or selecting maximal or minimal distances). The latent variable is populated as follows:

$$v_{isq}^{F-rad} = \begin{cases} 1, \; if \; d_q \in \{x_1^I \leq \cdots \leq x_S^I\}, \;\; and \; \sigma[|\boldsymbol{x}_{is}|: D_i] = \sigma[d_q: \{x_1^I, \dots, x_S^I\}] \\ \qquad 0, \qquad\qquad otherwise \end{cases} \quad , \qquad (1)$$

We are associating each target distance with the one distance from the model that has the same rank upon ordering all the model distances $D_i$. If $d_q$ is not a target distance (i.e., $d_q \notin \{x_1^I \leq \cdots \leq x_S^I\},$ ), we populate all those entries with a 0.

Lamina DamID: By using the information in the data $U^{LAM} \in \mathbb{R}^{(H)}$ and the population $\boldsymbol{X}^{(t)}$, we determine the latent variable $V^{LAM} \in \mathbb{R}^{(N \times S)}$. Let us consider the set $U_\lambda^{LAM} = U^{LAM}[U^{LAM} \geq \lambda]$, $\lambda$ being the already discussed lamina DamID probability threshold (*Methods*, **Extended Data Figures 1** and **2**). We then proceed as follows, by turning the contact probability into a frequency: in the set of all $2S$ sorted distances $d(\boldsymbol{x}_{is}, NE)$ to the nuclear envelope NE related to locus $I$, a lamina distance threshold $d_I^{act}$ is selected to be the $(2S \cdot e_I)th$ rank value. The latent variable $(i, i')$ slices are then populated by counting the number of all pooled distances that are larger than the corresponding activation distance, i.e.

$$v_{is}^{LAM} = \begin{cases} 1 \;\; if \; d(\boldsymbol{x}_{is}, NE) \geq d_I^{act} \\ 0 \quad otherwise \end{cases} \qquad\qquad i, i' \in I. \qquad (3)$$

**Bivariate data**, $I - J$ pair.

3D pairwise FISH By using the information in the data $\boldsymbol{M}^{F-dist} \in \mathbb{R}^{(H \times H \times Q)}$ and the population $\boldsymbol{X}^{(t)}$, we determine the latent variable $\boldsymbol{W}^{F-dist} \in \mathbb{R}^{(N \times N \times Q \times S)}$. The target distances for pair $I - J$ are given in the list $\{d_1, \dots, d_Q\}$ $(S \leq Q)$.

We compute all the pairwise distances $D_{ij} = \{|\boldsymbol{x}_{is} - \boldsymbol{x}_{js}|\}_{s=1, i \in I, j \in J}^{S}$ from structures in population $\boldsymbol{X}^{(t)}$, and sort them in ascending order: we obtain a sequence of ordered triplet indexes (diploid locus, diploid locus and structure):

$$\{(i_1, j_1, s_1), (i_2, j_2, s_2), \dots\} : \; |\boldsymbol{x}_{i_n s_n} - \boldsymbol{x}_{j_n s_n}| \leq |\boldsymbol{x}_{i_m s_m} - \boldsymbol{x}_{j_m s_m}|, \forall \, n \leq m$$

We subsample the list down to $S$ element (using either a stride or selecting maximal or minimal distances). Then, the latent variable is populated:

$$w_{ijsq}^{F-dist} = \begin{cases} 1, \ if \ d_q \in \{y_1^{IJ} \le \cdots \le y_S^{IJ}\}, \ and \ \sigma[|\boldsymbol{x}_{is} - \boldsymbol{x}_{js}|: D_{ij}] = \sigma[d_q: \{y_1^{IJ}, \dots, y_S^{IJ}\}] \\ 0, \qquad otherwise \end{cases} \quad (4)$$

We associated each target distance $y_{n=1,\dots,S}^{IJ}$ with the two diploid loci in the one structure in the population $\boldsymbol{X}^{(t)}$ that have the same rank in the distance distribution If $d_q$ is not a target distance (i.e., $d_q \notin \{y_1^{IJ} \le \cdots \le y_S^{IJ}\}$), then we populate that entry with a zero.

Ensemble Hi-C By using the information in the data $\boldsymbol{M}^{HiC} \in \mathbb{R}^{(H\times H)}$ and the population $\boldsymbol{X}^{(t)}$, we determine the latent variable $\boldsymbol{W}^{HiC} \in \mathbb{R}^{(N\times N\times S)}$. Let us consider the set $\boldsymbol{M}_{\theta}^{HiC} = \boldsymbol{M}^{HiC}[\boldsymbol{M}^{HiC} \ge \theta]$, $\theta$ being the Hi-C contact probability threshold (*Methods*, **Extended Data Figures 1** and **2**). If $I$ and $J$ are from the same chromosomes, only $(i,j)$ and $(i',j')$ combinations are considered; otherwise, all four distances $(i,j), (i,j'), (i',j)$ and $(i',j')$ are retained: overall, we have a set of 2S (4S) ordered pairwise distances, respectively. As before, we compute the activation distance $d_{IJ}^{act}$ as the $(2S \cdot a_{IJ})$-th $((4S \cdot a_{IJ}) - th)$ rank, respectively. The latent variable is populated by counting the number of all pooled distances that are shorter than the corresponding activation distance,

$$w_{ijs}^{HiC} = \begin{cases} 1, & if \ |\boldsymbol{x}_{is} - \boldsymbol{x}_{js}| \le d_{IJ}^{act} \\ 0, & otherwise \end{cases} \quad (i,i') \in I, (j,j') \in J. \quad (6)$$

i.e., a given contact is assigned to those structures whose participating domains are spatially fairly close already.

**Multivariate variable** Assume one spatial cluster is formed by $n$ haploid loci $\{I_1, \dots, I_n\}$. Using the information in the data $\boldsymbol{T}_n \in \mathbb{R}^{(H^n)}$ and the population $\boldsymbol{X}^{(t)}$, we determine the latent variable $\boldsymbol{R} \in \mathbb{R}^{(N^n \times S)}$.

The radius of gyration $R_g[I_1, I_2, \dots, I_n; s]$ of loci $\{I_1, \dots, I_n\}$ in population structure $s$ is an appropriate indicator and guides the search for the one structure that is already mostly compact. However, extensive search in the combinatorial space of all possible diploid cluster expansions grows exponentially with cluster size $n$. We circumvent this by assuming that loci mapping to the same chromosome number are also physically on the same copy. In order to determine which copy $h_\alpha$, a representative (haploid) locus $p_\alpha$ is randomly selected for each chromosome, $\alpha$. The set of optimal copy indexes is then determined from

$$\hat{h}_1, \dots, \hat{h}_n = \arg\min_{h_1,\dots,h_M} R_g^2[(I_1, h_1), \dots, (I_{n_{chr}}, h_{n_{chr}}); s]$$

as the radius of gyration among representative loci is being minimized in the parameter space of all combinations of copy indexes. If a cluster involves $n_{ch} \leq n$ chromosomes, there will be $n_{chr}$ representative loci and thus $2^{n_{chr}}$ combinations to explore, which can be enumerated easily than $2^n$. The loci forming the cluster in the diploid representation are then $(I_1, \hat{h}_1), \dots, (I_n, \hat{h}_n)$.

The following two chromosome toy model should help convey the idea. Assume we have a cluster formed by (1,2,20) loci from chromosomes (1,1,2) respectively $n = 3, n_{chr} = 2$, and assume both chromosomes come in two copies. Overall, there are six diploid loci:

| | $\boldsymbol{\alpha}$ | $\boldsymbol{h_1}$ | $\boldsymbol{h_2}$ |
|---|---|---|---|
| $\boldsymbol{I_1 = 1}$ | 1 | 1 | 201 |
| $\boldsymbol{I_2 = 2}$ | 1 | 2 | 202 |
| $\boldsymbol{I_3 = 20}$ | 2 | 20 | 220 |

We randomly select $I_1 = 2$ and $I_2 = 20$ as representative loci for each chromosome, and compute the radius of gyration in the space of the copy index h:

| $(I_1, h_{1)}$ | $(I_2, h_2)$ | $R_g$ |
|---|---|---|
| 2 | 20 | 102.6 |
| 2 | 220 | 33.2 |
| 202 | 20 | 155 |
| 202 | 220 | 300 |

The minimal radius of gyration is for $(h_1 = 1, h_2 = 2)$, so all loci from chromosome one are physically on the first copy, and loci from chromosome two are on the second, e.g. $(1, 2, 220)$.

Now, a given cluster would ideally be associated with the one structure in the pool that has the smallest radius of gyration; however, a very compact structure might be inconveniently overloaded with many clusters. In order to control this, we define the structure index probability distribution according to the following Gibbs factor:

$$P(s) = \frac{1}{Z} \exp\left[-\frac{R_g[s]}{kT} + \omega_s\right], \quad Z = \sum_s P(s) \qquad (7)$$

First, the average value of clusters per structure is introduced, $\langle n \rangle = S/n_{cl}$ (ratio between number of structures in the population and the total number of clusters to be assigned). Then, a penalization $\omega_s = \frac{\theta(occ_s - \langle n \rangle)}{\sqrt{\langle n \rangle}}$ is introduced, where $occ_s$ is the integer number of clusters already assigned to the current structure s, which we shall call occupancy, $\theta$ is the Heaviside step function. Note that a non-zero penalization is applied only when the current occupancy is larger than $\langle n \rangle$. Then,

$$r_{I_1,\ldots,I_n,s} = \begin{cases} 1, & if\ i = (c_1, \hat{h}_1), \ldots,\ \ and\ s = P^{-1}(s)] \\ 0 & otherwise \end{cases} \quad (8)$$

So that the index of the structure to which the cluster will be assigned is then drawn from the index distribution $P(s)$.

#### 2. A/M optimization: Modeling Step

Assignment provided a complete set of binary latent variables $v_{iqs}$, $w_{ijqs}$ and $r_{ijk\ldots s}$ which we would like to include in the genome models (see *Methods*). Each non-zero entry latent variable entry is modeled as an appropriate energy term (i.e., restraint), which is added to the full genome energy function. Each energy term involves one (univariate), two (bivariate) or more (multivariate) chromatin loci, and actively constrains their distances; each interaction is associated with a positive scalar (residual error) $\eta$ which monitors to what extent a given restraint is violated. Specifically, $\eta > 0.05$ indicates that a given distance in the model is off by 5% with respect to the input data: we will call that a violation.

##### Univariate variables.

3D radial FISH $v_{iqs}^{F-rad} = 1$ indicates that locus $i$ must be $d_q$ far from the center in structure $s$. We enforce that by a combination of truncated harmonic potentials with a tolerance $t$ (*Methods*, **Extended Data Figures 1** and **2**):

$$U^{F-rad}{}_{iqs} = \frac{k^{F-rad}}{2} \begin{cases} \left[|\boldsymbol{x}_{is}| - (d_q - t)\right]^2, if\ |\boldsymbol{x}_{is}| \le d_q - t \\ 0 \\ \left[|\boldsymbol{x}_{is}| - (d_q + t)\right]^2, if\ |\boldsymbol{x}_{is}| \ge d_q + t \end{cases} \quad (9)$$

The residual error ($0 \le \eta \le 1$) is defined as a ratio of distances:

$$\eta_{iqs}^{F-rad} = \begin{cases} \frac{|\boldsymbol{x}_{is}|}{d_q - t} - 1\,, if\ |\boldsymbol{x}_{is}| \le d_q - t \\ 0 \\ \frac{|\boldsymbol{x}_{is}|}{d_q + t} - 1, if\ |\boldsymbol{x}_{is}| \ge d_q + t \end{cases} \quad (10)$$

Lamina DamID $v_{is}^{LAM}$ indicates that locus $i$ is in contact with the lamina in structure $s$. So, we only apply an ellipsoidal lower harmonic bound (*Methods*, see also volumetric confinement):

$$U_{is}^{LAM} = \begin{cases} \frac{k_{DamID}}{2}\left(\frac{1}{\sqrt{\kappa_i(r_0,c_r)}} - 1\right)^2 |\boldsymbol{x}_{is}|^2, & if\ |\kappa_{is}| \le 1 \\ 0 & otherwise \end{cases}, \quad (11)$$

$$\kappa_{is}(r, c_r) = \frac{x_{is}^2}{[a(1-c_r)-r_0]^{\,2}} + \frac{y_{is}^2}{[b(1-c_r)-r_0]^{\,2}} + \frac{z_{is}^2}{[b(1-c_r)-r_0]^{\,2}} \quad (12)$$

Where we define a contact shell as the volume $\Omega_0/\Omega(a(1-c_r)-r_0, b(1-c_r)-r_0, c(1-c_r)-r_0)$ between two concentric ellipses, $c_r$ being a positive scalar (this is our approximation to the lamina contact threshold $l^{contact}$, see *Methods*): $c_r = 0$ implies that contact is only formed when the genome locus is physically in contact with the envelope.

The residual error is here evaluated as the ratio between the distance of a locus to the lamina and its radial distance from the center:

$$\eta_{is}^{LAM} = \begin{cases} (1-\sqrt{\kappa_{is}}), if\ \kappa_{is} < 1 \\ 0 \end{cases} \quad (13)$$

and is non-zero when the locus is not in the outer contact shell.

**Bivariate variables**.

3D pairwise FISH $w_{ijqs}^{F-dist}$ indicates that the pairwise distance of loci $i$ and $j$ in structure $s$ is equal to $d_q$. We then use the same combination of potentials which now act on the mutual distance, with tolerance $t$ (*Methods*, **Extended Data Figures 1** and **2**):

$$U_{ijqs}^{F-dist} = \frac{k^{F-dist}}{2}\begin{cases} \left[|\boldsymbol{x}_{ijs}| - (d_q - t)\right]^2, if\ |\boldsymbol{x}_{ijs}| \le d_q - t \\ 0, \quad if\ d_q - t \le |\boldsymbol{x}_{ijs}| \le d_q + t \\ \left[|\boldsymbol{x}_{ijs}| - (d_q + t)\right]^2, if\ |\boldsymbol{x}_{ijs}| \ge d_q + t \end{cases} \quad (14)$$

The residual error ($0 \le \eta \le\ 1$) is defined as:

$$\eta_{ijqs}^{F-dist} = \begin{cases} \frac{|\boldsymbol{x}_{ijs}|}{d_q - t} - 1\,, if\ |\boldsymbol{x}_{isj}| \le d_q - t \\ 0, \quad if\ d_q - t \le |\boldsymbol{x}_{ijs}| \le d_q + t \\ \frac{|\boldsymbol{x}_{ijs}|}{d_q + t} - 1, if\ |\boldsymbol{x}_{ijs}| \ge d_q + t \end{cases} \quad (15)$$

Ensemble Hi-C The reduced variable $w_{ijs}^{HiC}$ indicates that $i$ and $j$ form a contact in structure $s$. We then only apply the upper harmonic bound ($C$ labels the contact distance)

$$U_{ijs}^{HiC} = \begin{cases} \frac{k_{HiC}}{2}\left[\left|\boldsymbol{x}_{ijs}\right| - l^C\right]^2, if \left|\boldsymbol{x}_{ijs}\right| \geq l^C \\ 0 \end{cases}, \quad l^C = 4r_0. \ (16)$$

$$\eta_{ijs}^{HiC} = \begin{cases} \frac{\left|x_{ijs}\right|}{C} - 1, if \left|\boldsymbol{x}_{ijs}\right| \geq l^C \\ 0 \end{cases} \qquad (17)$$

**Multivariate variables**. $r_{i_1,\dots,i_n,s} = 1$ indicates that loci $i_1, \dots, i_n$ form a cluster in structure $s$. We introduce a massless particle (centroid, no excluded volume effect) in structure $s$ located in the cluster geometric center, $\boldsymbol{x}_G = \frac{1}{n}\sum_{p\in\{i_1,\dots,i_n\}} \boldsymbol{x}_{ps}$. We then introduce the following spatial restraint:

$$U_{i,\dots,i_{n,s}}^{SPRITE} = \begin{cases} \frac{k_{SPRITE}}{2}\sum_{p\in\{i,\dots,i_n\}}\left[\left|\boldsymbol{x}_{ps} - \boldsymbol{x}_G\right| - (r_n - r_0)\right]^2 & if \left|\boldsymbol{x}_p - \boldsymbol{x}_G\right| \geq r_n - r_0 \\ 0 & otherwise \end{cases},(18)$$

where $r_n = r\sqrt[3]{n/\rho}$ indicates the chosen radius associated with a cluster of $n$ loci and $\rho$ volumetric density (*Methods*, **Extended Data Figures 1** and **2**), and loci are harmonically restrained to the centroid in order to promote geometric integrity. Each of the terms in the summation are associate with a residual error Eq. (17).

$$\eta_{i,\dots,i_{n,s}} = \{\eta_p\}_{p=\in\{i_1,\dots,i_{n,s}\}}, \qquad \eta_p = \begin{cases} \frac{|x_p - x_G|}{r_n - r_0} - 1, & if \left|x_p - x_G\right| \geq r_n - r_0 \\ 0 & otherwise \end{cases} (19)$$

**Polymer restraints** Each genome chromosome is represented as a free chain of 200k base-pair resolution coarse-grained loci, subjected to connectivity and steric restraints, i.e.

$$U_{polymer} = U_{steric} + U_{chain} = \sum_{i\neq j} A_{ij}^{chain}\left[1 + \cos\left(\frac{\pi\left|\boldsymbol{x}_{is} - \boldsymbol{x}_{js}\right|}{R_C}\right)\right] + \sum_{i,i+1}\frac{K_b}{2}\left[\left|\boldsymbol{x}_{is} - \boldsymbol{x}_{js}\right| - C\right]^2$$

$$R_c = 2r_0, \qquad A_{ij}^{chain} = \left(\frac{2r_0}{\pi}\right)^2,$$

Steric interactions are modeled using a soft potential with cutoff $R_c$ and interaction amplitude $A_{ij}^{chain}$ and are bivariate, i.e. act between two loci. The interconnectivity term between neighboring loci (within the same chain) is a harmonic upper interaction (i.e., only acts when the distance exceeds the contact distance $l^C$) of elastic constant $K_b$. The associated residual errors read:

$$\eta_{steric,s} = \begin{cases} 0, & if\ |\boldsymbol{x}_{ijs}| \geq R_c \\ \frac{|\boldsymbol{x}_{ijs}|}{R_c} - 1, & otherwise \end{cases}, \qquad \eta_{chain,s} = \begin{cases} \frac{|\boldsymbol{x}_{ijs}|}{l^C} - 1, if\ |\boldsymbol{x}_{ijs}| \geq l^C \\ 0, \quad otherwise \end{cases}$$

**Nuclear confinement** A lamina DamID restraint Eqs. (11)-(12) with a negative (attractive) elastic constant efficiently models the effects of the volume confinement of the nuclear lamina (*Methods*):

$$U_{vol,s} = \begin{cases} -\frac{|k_{DamID}|}{2}\left(\frac{1}{\sqrt{\kappa_{is}(r,0)}} - 1\right)^2 |\boldsymbol{x}_{is}|^2, & if\ |\kappa_i| \geq 1 \\ 0 & otherwise \end{cases}$$

In this specific case, the contact range $c_r$ is set to zero: the restraint is applied any time the geometric center of a locus is outside of the lamina. The residual error is:

$$\eta_{vol,s} = \begin{cases} \left(1 - \sqrt{1/k_{is}}\right), & if\ \ \kappa_i > 1 \\ 0 & otherwise \end{cases}$$

#### 3. Synthetic data simulations

Simulated Hi-C, lamina DamID and FISH (both radial and pairwise) data were extracted from the ground truth population as detailed in the *Methods* section. In particular, FISH (haploid) probes $I$ were selected randomly across all chromosomes, FISH pairs $(I,J)$ were selected by first downsampling the 15453 haploid loci using a uniform stride of 5, building all the pairs out of those, and then sampling randomly across this data set. A couple of technical remarks are in order. First, the same exact set of simulation parameters was used to generate all populations, in order to ensure consistency: eventually, all simulated populations satisfy more than 99.995% of the imposed restraints.

Second, data are incorporated into a population in a fashion similar to the steps adopted for real data, by resorting to decreasing optimization thresholds (see also **Extended Data Figures 1-2**). In particular, non-zero final threshold values are used here too, which represent the highest accuracy we are confident the models could achieve. This reproduces the standard scenario where data are only known within a confidence range, and probability values that are too small could be misconstrued as noise or a perturbation.

#### 4. Iterative contact refinement step

Since loci that are not expected to form a contact are not explicitly being restrained from doing that, excess contacts cannot be prevented, which are shown to lead to excessively compact structures that cannot be relaxed easily. A heuristic method has been put it place to compensate for such an effect, by allowing to assess the actual portion of expected contacts that are available for allocation, which then affects the way the activation distance $d_{act}^{IJ}$ is computed in Hi-C and DamID assignment steps. The empirical procedure relies on the predicament that when a population expresses more contacts than it should, we reduce the assignment probability: a lower probability (with fewer restraints) should be equally effective.

Assume the expected number of contacts $Sp_{ij}^{input}$ is the sum of a number of effective contacts that are actually imposed, $N_{eff}$, and a number of incidental contacts (that are also expressed but not imposed), $N_{inc}$:

$$Sp_{ij}^{input} = N_{eff} + N_{inc} = N_{eff} + \eta(S - N_{eff})$$

The number of incidental contacts is here expressed as a fraction of the number of non-applied (non-enforced) contacts. The latter term can originate from cooperative effects, which automatically bring loci closer without an explicit bonding term operating. We can solve for probability $p_0 = N_{eff}/S$: $p_0 = \frac{p^{input} - \eta}{1-\eta}$. This is an effective probability which controls the number of restraints to be enforced. Now, we need an estimate for the scalar $\eta$.

Let us to compare the factual number of contacts in the population (expressed by the tensor $A_{ij}^{X} = \sum_{s=1}^{S} \boldsymbol{W}_{ijs}^{(k-1),\mathrm{X}}$) with the predicted number of contacts from the previous assignment step ($A_{ij}^{assig} = \sum_{s=1}^{S} \boldsymbol{W}_{ijs}^{(k-1),\mathrm{assig}}$):

$$\sum_{s=1}^{S} \boldsymbol{W}_{ijs}^{(k-1),\mathrm{X}} = \sum_{s=1}^{S} \boldsymbol{W}_{ijs}^{(k-1),\mathrm{assig}} + \left(S - \sum_{s-1}^{S} \boldsymbol{W}_{ijs}^{(k-1),\mathrm{assig}}\right)\eta_{ij}$$

We can solve for $\eta_{ij}$:

$$\eta_{ij} = \frac{\sum_{s=1}^{S} \boldsymbol{W}_{ijs}^{(k-1),\mathrm{X}} - \sum_{s=1}^{S} \boldsymbol{W}_{ijs}^{(k-1),\mathrm{X}}}{S - \sum_{s=1}^{S} \boldsymbol{W}_{ijs}^{(k-1),\mathrm{assig}}} = \frac{A_{ij}^{X} - A_{ij}^{assig}}{1 - A_{ij}^{assig}},$$

which can then be plugged into the previous equation to find a corrected assignment probability $p_0 = \frac{p^{input} - \eta}{1-\eta}$, which is then used to update the activation distance $d_{ij}^{act}$. Please note that the correction is only implemented if there is a contact excess.

#### 5. HFFc6 experimental data pre-processing

##### Ensemble Hi-C data[1]

4DN portal identifier: 4DNES2R6PUEK

We used *in situ* Hi-C datasets from HFFc6 cell line (reference genome hg38)[1]. Similar to the protocol by ref.[2], low bin sequence coverage 3% regions were discarded during the normalization process. For data normalization, we adopted the same KR normalization method used in ref.[3], leading to a normalized contact frequency matrix $\boldsymbol{F} = (f_{ij})_{K\times K}$ at 20 kb resolution. We then generated probability matrix at 200 kb level as our further input for our algorithm using the following way:

The contact between 2 large domains is defined by the closest part of each other. We defined a mapping $b(i)$ as the set of all bins in matrix $\boldsymbol{P}$ that belongs to 200-kb region $i$. Then the domain-level matrix $A = (a_{ij})_{N\times N}$ was calculated as:

$$a_{ij} = mean(\ top10\% < \{p_{\alpha\beta}\colon\ \alpha \in b(i), \beta \in b(j)\} >)$$

In the case that some contacts are extremely higher than the surrounding contacts, these contacts were identified as outliers by $\{p\colon p > \mu + 1.5IQR\ \}$, where $p \in \{p_{\alpha\beta}\colon \alpha \in b(i), \beta \in b(j)\}$ and $\mu =$

$mean\{p_{\alpha\beta}: \alpha \in b(i), \beta \in b(j)\}$. The IQR refers to the interquartile range of $\{p_{\alpha\beta}\}$. These outliers were excluded from the calculation.

Next, we converted the contact frequencies in the 200-kb matrix to contact probabilities by scaling the frequencies by a normalization factor, $f^{max}$, which is chosen to represent the contact frequency value at which two domains have a 100% probability to form a contact. The 200-kb contact probability matrix $\boldsymbol{P} = (p_{ij})_{K\times K}$ was calculated as $p_{ij} = \min\left(\frac{f_{ij}}{f^{max}}, 1\right)$, where $p_{ij}$ and $f_{ij}$ are the contact probability and frequency values, respectively. We set the value of $f^{max}$ so that the average contact probability sum of a 200-kb region is ~24, which, based on our experience, is the average number of contacts a domain has at saturation (where no more contact restraints can be satisfied)[3].

After obtaining the contact probability matrix at 200-kb resolution, we identified bins that have spurious inter-chromosomal interaction probabilities (higher than 0.2), and removed the corresponding bins in the 20-kb raw matrix, and repeated the KR normalization, and regenerated 200-kb contact probability matrix where no inter-chromosomal contact probability is higher than 0.2. Finally, we set contact probabilities between the consecutive domains as well as between domains up to 1 Mb distance in the gap regions to 1 in order to maintain the chain integrity.

It is possible that the elongated HFF shape may require a different normalization of the raw Hi-C data when mapping onto the 200kb resolution than what has been successfully used with the GM cell line. In order to avoid biases, we performed extensive benchmarking by generating multiple populations with different setups and eventually assessing the quality of orthogonal quantities prediction. Our analysis revealed that the best setup requires scaling up all the inter-chromosomal interactions by a factor of 1.5 and including all those that are more frequent than 0.4% in the optimization. We observed that an inappropriate scaling can still be circumvented by adding other data modalities to the population, which confirmed the results from the synthetic calculation section.

**Lamina DamID[4]**

4DN portal identifier: 4DNESXZ4FW4T

Mapping data to 200-kb resolution

Each region (5-kb) in the Lamina DamID data was mapped to 200-kb regions with an overlap of 50% or higher. After mapping, the signals of the multiple 5-kb regions mapped to each 200-kb region were averaged (we first took the inverse log2 of the signals, then averaged and took the log2 of the average value).

Estimating lamina contact probabilities from Lamina DamID signals

We would like to convert the experimental lamina DamID signals into an array of lamina contact probabilities $e_I$. First, we assume that only chromatin regions with a raw signal larger than 1 ($sig_I \geq 1.0$, Dam-LaminB1/Dam ratio), which represent 54% of the whole genome for HFF, would have a lamina contact probability larger than 0, therefore we set all signals lower than 1 to 0. Next, we utilize from the single-cell lamin DamID data[5] to relate raw signals to lamin contact probabilities. We first calculate the average contact probability observed in the single-cell lamin DamID data[5] and use that as a conversion factor as follows:

$$e_I = \frac{sig_I}{\overline{sig_I}} mean(Contact\ probabilities\ from\ KIND\ et\ al.)$$

where $\overline{sig}$ is the genome-wide average of all signals. This way, we have $\overline{e_I}$, the average of our estimated lamina contact probabilities is equal to the average calculated from the single cell lamina DamID data[5] .With this calculation, we obtain an estimated DNA content near the nuclear envelope (NE) to be 28%. We also tested different values of DNA content at the NE between 15 – 40 % and carried out extensive parameter search to identify the optimal value: Each setup was used to generate a Hi-C + lamina DamID population, and lamina DamID, SON TSA-seqs and 3D HIPMAp FISH data were computed and compared with available experimental data (see below for preprocessing of those data). A cumulative ranking was computed and suggested that using a percentage of ~25% performs best: that is the setup we used for all real data lamina DamID calculations discussed in the main text.

**3D HIPMap data**[6]

4DN portal identifiers: https://data.4dnucleome.org/publications/80007b23-7748-4492-9e49-c38400acbe60/

We looked into the summary table accompanying the raw data and only retained those pairs passing the accuracy test[6]. Each FISH probe was mapped onto a 200kb resolution in our model. For each pair, all distances provided in the raw data were extracted and ordered to obtain a discrete cumulative distribution of distances. We used a polynomial Savoy-Golay filter[7] for interpolating each cumulative distribution and drew 1000 uniformly sampled distance values from that. The resulting set of 1000 distances per distribution are the target distances that are then restrained in the simulation. Specifically, those distances we use as minimal FISH distances (see *Assignment*). In the simulations we only included those pairs separated by at least 10Mb genomic distance (resulting to 51 pairs).

**DNA SPRITE data**[8]

4DN portal identifier: 4DNESJYGTI8S

We first filtered out mitochondrial chromatin and outliers. We then mapped colocalized loci to 200kb resolution and selected fully inter-chromosomal clusters involving different loci only, up to 6 chromosomes (overall, 6617 clusters). We generated the appropriate family of tensors $\boldsymbol{T}_n = (t_{I_1,\dots I_n})$ (see *Methods*) and used it as input to IGM calculations.

DNA SPRITE data for HFFc6 was kindly provided by the Guttman lab at CALTECH, and are not yet publicly accessible.

**SON TSA-seq**[9]

4DN portal identifier: 4DNES85R9TIB

Each region in the TSA-seq data was mapped to 200-kb regions with an overlap of 50% or higher. After mapping, each 200-kb region had multiple TSA-seq regions, therefore the signals mapped to each 200-kb region were averaged (we first took the inverse log2 of the signals, then averaged and took the log2 of the average value).
